# Control of topoisomerase II activity and chemotherapeutic inhibition by TCA cycle metabolites

**DOI:** 10.1101/2021.01.22.427624

**Authors:** Joyce H. Lee, Eric P. Mosher, Young-Sam Lee, Namandjé N. Bumpus, James M. Berger

## Abstract

Topoisomerase II (topo II) is essential for disentangling newly replicated chromosomes. DNA unlinking involves the physical passage of one DNA duplex through another and depends on the transient formation of double-strand DNA breaks, a step exploited by frontline chemotherapeutics to kill cancer cells. Although anti-topo II drugs are efficacious, they also elicit cytotoxic side effects in normal cells; insights into how topo II is regulated in different cellular contexts is essential to improve their targeted use. Using chemical fractionation and mass spectrometry, we have discovered that topo II is subject to metabolic control through the TCA cycle. We show that TCA metabolites stimulate topo II activity *in vitro* and that levels of TCA flux modulate cellular sensitivity to anti-topo II drugs *in vivo*. Our works reveals an unanticipated connection between the control of DNA topology and cellular metabolism, a finding with important ramifications for the clinical use of anti-topo II therapies.

## INTRODUCTION

The complement of chromosomes in a single human cell, laid out end-to-end, is ∼10,000 times longer than the diameter of the nucleus in which they reside. Efficient packaging, access, and duplication of this genetic material depends on the proper control of DNA superstructure. Some of the largest physical rearrangements of the genome occur during DNA replication and mitosis as sister chromatids are synthesized, condensed, and segregated into two daughter cells (Barrington et al., 2017). The double-helical structure of DNA presents a challenge to these transformations, resulting in superhelical intertwinings and chromosomal entanglements (catenanes) (Peter et al., 1998; Postow et al., 2001; Sundin and Varshavsky, 1981). Enzymes known as topoisomerases are required by cells to resolve topological stresses in DNA; of the two major classes, only type II topoisomerases are able to unlink tangled double-stranded segments to facilitate chromosome partitioning prior to cell division (Vos et al., 2011).

Beyond supporting supercoiling homeostasis and promoting DNA unlinking, mounting evidence suggests that topo II has additional roles in regulating chromatin structure throughout the eukaryotic cell cycle (Lee and Berger, 2019). Although most DNA regions are decatenated by the end of replication in S phase (Charbin et al., 2014; Lucas et al., 2001), the maintenance of some catenated regions, mostly in regions with repetitive sequences, appears to be important for proper chromosome condensation and sister chromatid cohesion (Bauer et al., 2012; Daniloski et al., 2019). Loss or gain of topo II function has been shown to disturb the precisely balanced catenation state of the genome and lead to detrimental effects on chromosome condensation (Cuvier and Hirano, 2003; Samejima et al., 2012; Uemura et al., 1987). In vertebrates, the cell cycle-dependent expression of topo IIα, one of two topo II isoforms found in such organisms, provides one means of calibrating levels of topoisomerase activity (Heck et al., 1988; Kimura et al., 1994; Woessner et al., 1991); however, invertebrates and lower-order eukaryotes (such as yeast and *Drosophila*) express only a single isoform of topo II for which cell cycle-dependent expression has not been observed (Eser et al., 2011; Goto and Wang, 1984; Spellman et al., 1998; Whalen et al., 1991). Post-translational modifications such as SUMOylation ubiquitylation, and phosphorylation – the majority of which map to an unconserved, species-specific C-terminal domain of topo II – have been reported to regulate enzyme stability, localization and activity, but how these marks exert such functions is poorly understood (Lee and Berger, 2019). Collectively, these and other observations demonstrate that there is requirement for precise regulation of topo II activity. However, whether there exist fundamental regulatory mechanisms that are conserved throughout eukaryotes has not been established.

Because topo II is essential for cell proliferation, it is a potent target for many chemotherapeutics (Nitiss, 2009). A special class of topo II-targeting drugs, categorized as topoisomerase ‘poisons,’ triggers cell death by inducing topo II to generate cytotoxic DNA damage (Pommier et al., 2010; Wu et al., 2011). The major limitation of topoisomerase poisons is their potential to generate DNA damage in noncancerous cells that can lead to serious toxic side effects and therapy-related neoplasias (Felix, 1998; Turcotte et al., 2018). A greater understanding of the cellular mechanisms that diminish or enhance topo II activity levels is necessary to more optimally match topoisomerase-targeted therapies to specific cancer types and to design combinatorial strategies that might selectively enhance the potency of anti-topoisomerase drugs in cancer cells.

In a prior study, we found that a highly-conserved pocket in the ATPase domain of topo II that binds to ICRF-187, a clinically-approved drug, also associates with a natural, plant-derived antagonist of topo II known as resveratrol (Lee et al., 2017). This observation led us to wonder whether other small molecules found natively in cells might also modulate topo II function. To test this idea, we prepared crude metabolite extracts from *S. cerevisiae* and analyzed them for activity against topo II *in vitro*. We discovered that these extracts were capable of stimulating topo II activity, an effect that has not been observed previously for other agents that act on topo II. Using biochemical fractionation and LC-MS/MS analysis, we determined that di- and tri-carboxylate TCA cycle metabolites were responsible for the stimulatory effect, and that these compounds increase the efficiency of the topo II strand passage reaction. By monitoring the sensitivity of yeast to different classes of topo II inhibitors under different metabolic states, we further found that TCA cycle flux can directly influence topo II activity *in vivo.* Collectively, our results show that natural small molecules produced by metabolic processes of the cell can directly modulate topo II function and that manipulation of these processes affects the efficacy of clinically-approved topo II-targeting drugs. These findings in turn reveal an unanticipated link between cellular metabolism and the regulation of a key central dogma process, a discovery that provides new directions for improving the safety and efficacy of commonly used anti-topo II chemotherapies.

## RESULTS

### Small-molecule metabolites from *S. cerevisiae* stimulate topoisomerase II activity *in vitro*

To search for natural products that might regulate topo II, we first tested whether crude *S. cerevisiae* metabolite extracts could impact enzyme activity *in vitro*. Metabolic extracts were prepared from yeast cultures grown in minimal media without ammonium sulfate, as sulfate ions were found to enhance topo II supercoil relaxation activity (**Figure S1A**) (all other media components were confirmed to have no significant effect on topo II activity (**Figure S1B**)). Small-molecule extracts were taken from log-phase cultures to capture metabolic profiles representative of actively dividing cells in all cell cycle stages (**Figure 1A**). A sample of the leftover spent media was also lyophilized as a control for media components and any secreted compounds.

**Figure 1.**
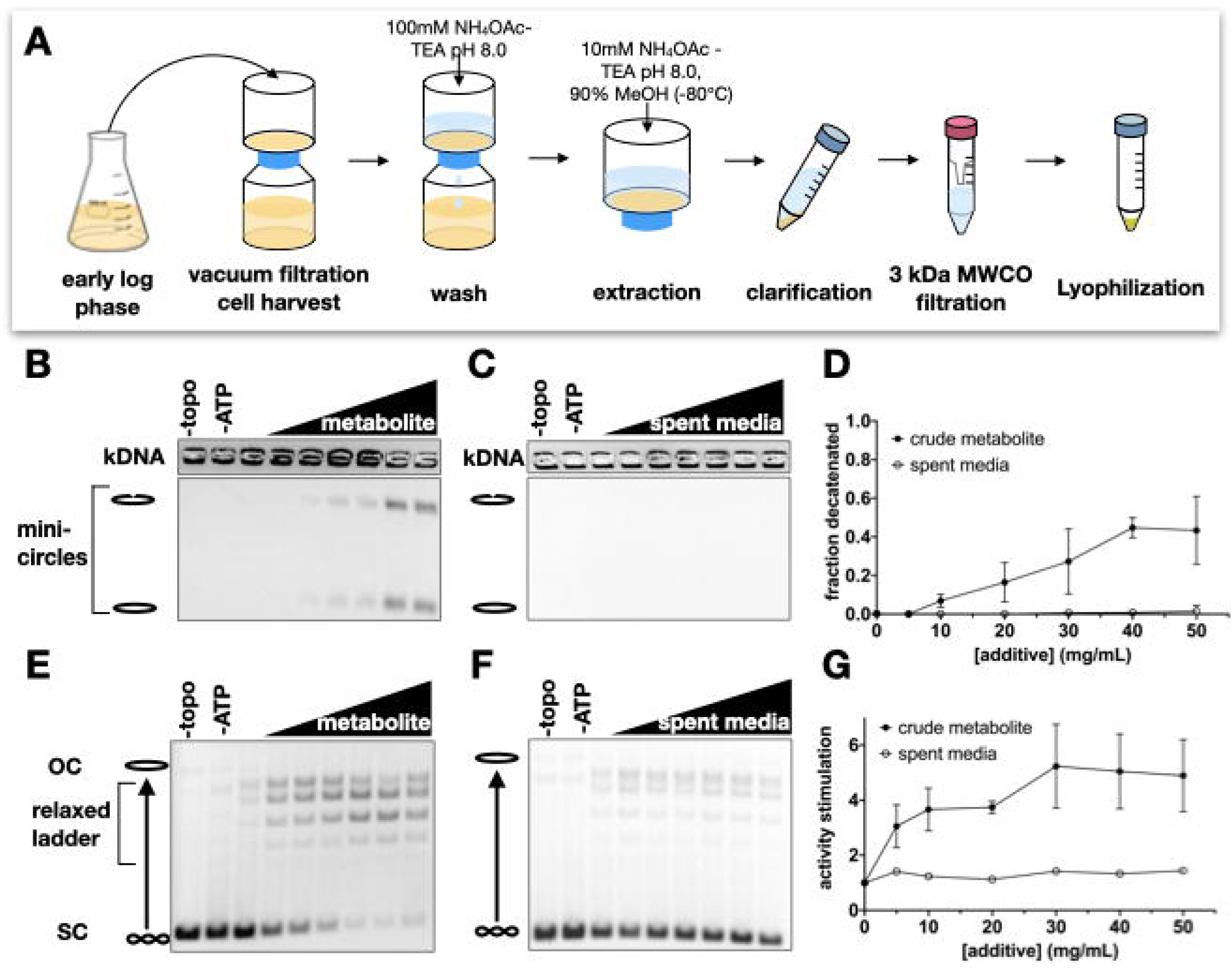
Crude metabolite extracts from log-phase yeast stimulate *Sc*Top2 strand passage activity. (A) Schematic of metabolite extraction procedure from yeast cultures. (B-C) Representative gels of kinetoplast DNA (kDNA) decatenation assays with metabolite extracts and spent media control samples. Bands representing nicked minicircles and closed minicircles are indicated to the left of the gels. No enzyme (-topo) and no ATP (-ATP) negative controls show the starting substrate. Lyophilized samples of the metabolite extract and spent media were titrated from 0 to 50 mg/ml in 10 mg/ml increments. (D) Decatenation assay data represented as mean ± SD of n=3 independent experiments. (E-F) Representative gels of supercoil relaxation assays with metabolite extracts and spent media control samples. The bands representing unrelaxed substrate (SC), the relaxed topoisomer distribution, and nicked/open circle (OC) plasmids are indicated to the left of the gels. No enzyme (-topo) and no ATP (-ATP) negative controls show the starting substrate. Lyophilized samples of the metabolite extract and spent media were titrated from 0 to 50 mg/ml in 10 mg/ml increments. (G) Supercoil relaxation assay data represented as mean ± SD of n=3 independent experiments.

We next assessed the effects of the crude metabolite extract and the spent media control on topo II using decatenation assays. Purified *S. cerevisiae* topo II (*Sc*Top2) was incubated with kinetoplast DNA (kDNA) – a highly interlocked network of small circular DNAs present in certain protozoa – and ATP to catalyze strand passage. All experiments were conducted with excess ATP (5 mM) to eliminate any response that might have been elicited from the presence of nucleotide in the extracts. Although the spent media sample had little to no effect on DNA unlinking by *Sc*Top2, the crude metabolite extract stimulated decatenation activity in a dose-dependent manner (**Figures 1B-D**). Similar to results obtained from decatenation assays, the crude extract also stimulated DNA supercoil relaxation by *Sc*Top2, as evidenced by the conversion of a negatively supercoiled plasmid substrate into a distribution of relaxed topoisomers (**Figures 1E-G**).

### Biochemical characterization of active compounds from yeast metabolite extracts

Upon discovering a stimulatory activity in crude extracts, we set out to define the chemical properties of the active agent(s). Liquid-liquid extraction with butanol was first conducted under both acidic and basic conditions. In both cases, the stimulatory activity was recovered in the aqueous phase, indicating that the agent was highly polar (**Figures S2A and S2B**). Based on this finding and the nucleotide-binding capabilities of topo II, we surmised that the factor might contain phosphate groups or phosphodiester bonds. However, the stimulatory activity was resistant to treatment with Antarctic phosphatase or snake venom phosphodiesterase, suggesting that neither phosphates nor phosphodiester bonds were present in the stimulatory metabolite (**Figures S2C and S2D**).

To further characterize the active compound and prepare samples for mass spectrometry analysis, we developed a purification protocol for the agent (**Figure 2A**). Solid-phase extraction (SPE) was used as a first-pass, bulk-fractionation method to remove contaminants and enrich the stimulatory activity roughly two-fold (**Figure 2B**). The active fraction from the SPE step was then further fractionated using reverse-phase HPLC, followed by normal phase HPLC (**Figures 2C-F**). Interestingly, this procedure not only enriched the stimulatory agent, but also revealed two separate fractions (the gray and pink fractions in **Figures 2D and 2F**) that inhibited DNA supercoil relaxation by *Sc*Top2 (**Figure 2F**); these latter activities appear to have been masked by the stimulatory activity in previous purification steps. The appearance of these distinct activities indicates that cells contain more than one class of small molecules that can act on topo II.

**Figure 2.**
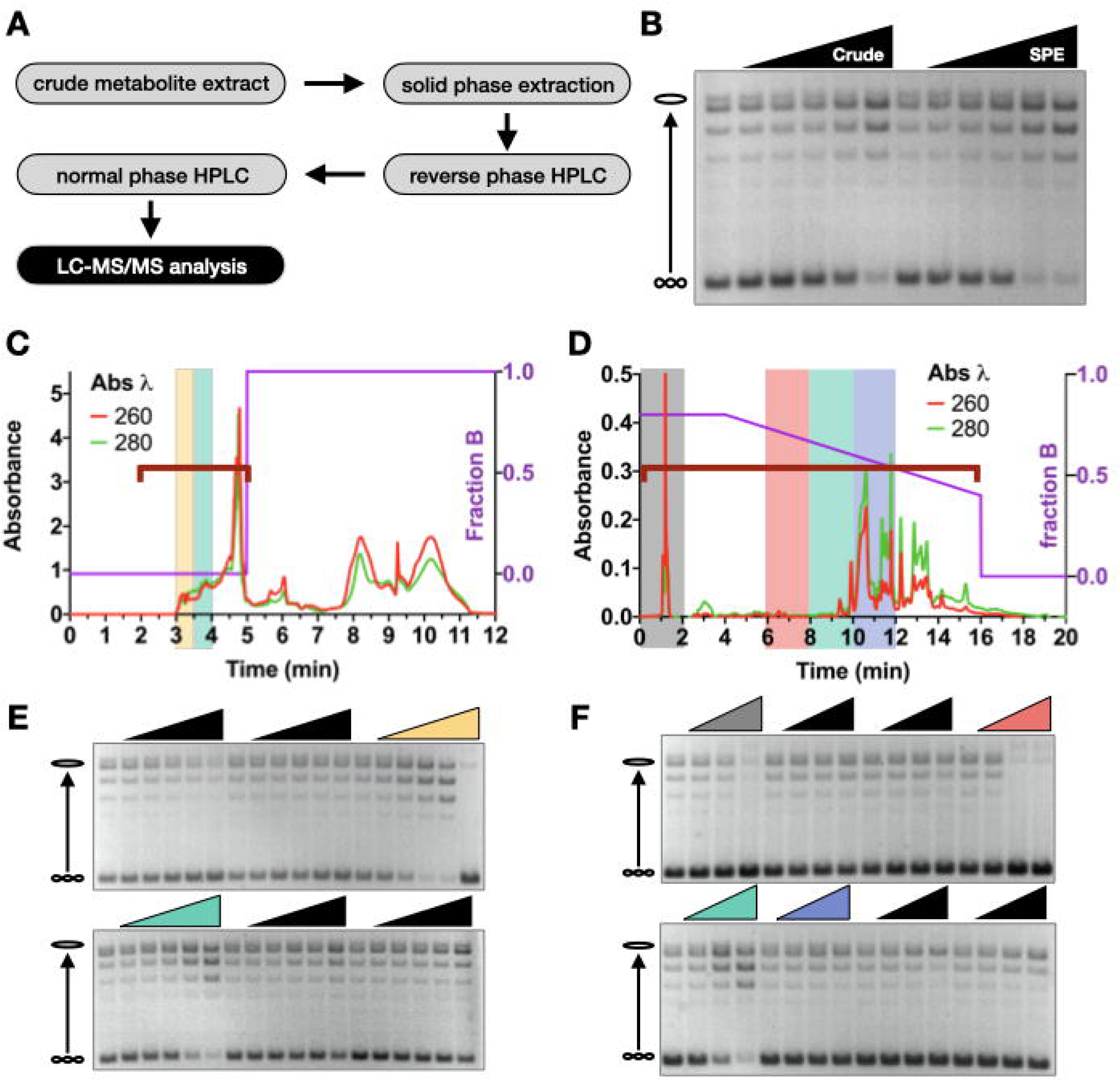
Purification of stimulatory metabolites from crude metabolite extracts. (A) Schematic of purification steps to prepare samples with enriched stimulatory activity for LC-MS/MS analysis. (B) Supercoil relaxation assay with metabolite samples before (‘Crude’) and after solid phase extraction (‘SPE’). Crude and SPE purified metabolite extracts were added from 0 to 40 mg/ml in two-fold increments (0, 2.5, 5, 10, 20, 40 mg/ml). (C-D) Chromatograms of reverse phase (C) and normal phase (D) HPLC purification runs. The HPLC method is depicted by the purple line showing fraction of phase B as indicated by the right y-axis. Red and green traces show absorbance values at 260 nm and 280 nm wavelengths respectively as indicated on the left y-axis. (E-F) Supercoil relaxation assays of reverse phase fractions (E) and normal phase fractions (F) indicated by the red brackets in (C) and (D). Fractions of interest are highlighted by corresponding colors in the chromatograms and relaxation assay gels. Lyophilized material from each fraction was solubilized in equal volumes of 12.5% DMSO and titrated down from the maximum possible concentration in two-fold dilution steps

### Identification of topo II-stimulating metabolites

To identify the stimulatory agent, the stimulatory fraction (green in **Figures 2D and 2F**) and flanking fractions (pink and blue in **Figures 2D and 2F**) from the last HPLC purification step were analyzed by LC-MS/MS. Spectra were first collected from a pooled sample to generate a peak-identification library that was representative of all metabolites present in the three samples. Data were processed in Compound Discoverer^TM^ by referencing mass spectral databases of natural products (**Figure 3A**). Next, the samples were analyzed individually and spectra from each sample were indexed based on the pooled library. By comparing peak intensities across samples, we generated a candidate list of compounds that were enriched in the stimulatory fraction as compared to its neighboring fractions (**Figure 3B, Table S1**).

**Figure 3.**
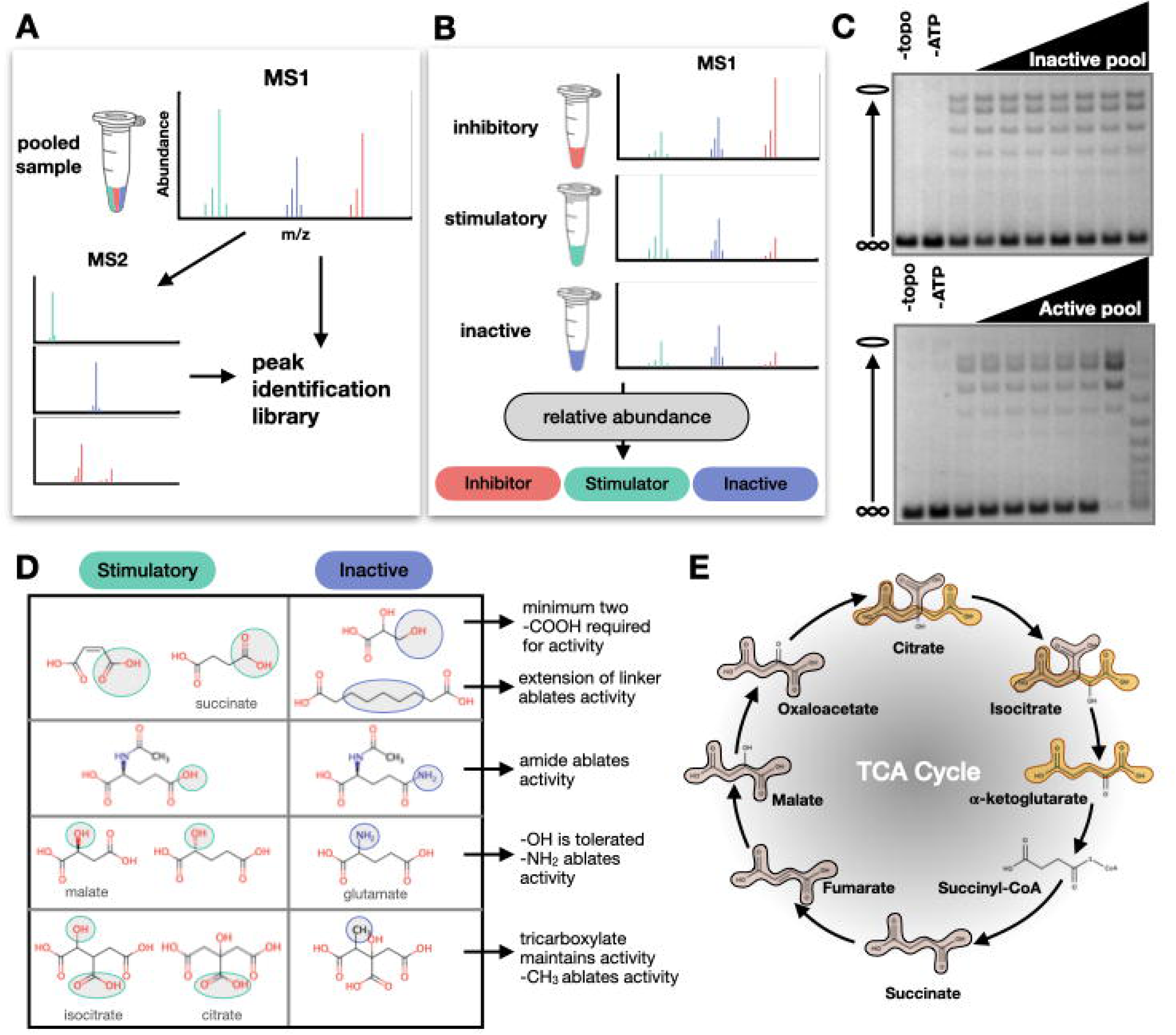
Identification and structure activity relationship (SAR) analysis of stimulatory compounds. (A) Schematic depicting how three sequential fractions (indicated by pink, green, and blue in Figure 2) were pooled and analyzed by LC-MS/MS to generate an ion peak identification library. (B) Schematic depicting how different samples (colored as pink, green, and blue) were analyzed individually. Intensities of metabolite peaks were compared across the three samples to determine relative abundance of each ion. Ions that were enriched in the stimulatory fraction (green) were added to a candidate list for stimulatory compounds (see **Table S1**) (C) Example supercoil relaxation assays from testing an inactive and an active pool of five candidate compounds. Candidate pools were prepared from commercial compounds. The concentration of each candidate compound was titrated from 0 to 20 mg/ml in two-fold steps (0, 0.31, 0.63, 1.25, 2.5, 5, 10, 20 mg/ml). (D) SAR analysis comparing stimulatory and inactive compounds identified from candidate pools. Chemical properties that are necessary for activity (green circles) and chemical changes that ablate activity (blue circles) are highlighted in gray. (E) TCA cycle metabolites. Succinic acid (tan) and glutaric acid (gold) motifs are highlighted.

Thirty-two candidate compounds were initially tested in eight different pools for stimulatory activity in supercoil relaxation assays. Although most pools showed no effect on topo II, some pools did impact enzymatic activity (**Figure 3C**); compounds from pools displaying activity were then tested individually. Several compounds from the initial candidate list showed an ability to stimulate DNA supercoil relaxation by *Sc*Top2. To better understand the chemical properties necessary for this stimulation, we compared the chemical structures of active compounds to those of inactive compounds with similar features (**Figure 3D**). These structure-activity relationship (SAR) analyses revealed that short-chain dicarboxylic acids were able to stimulate topo II activity.

Compounds with an additional hydroxyl group on the methylene chain maintained activity, but an amine or a methyl group in the same position ablated the stimulatory effect. Tricarboxylate compounds were also able to stimulate topo II activity, whereas extension of the connecting linker beyond three carbons ablated compound activity (**Figure 3D**). Together, these tolerable and intolerable chemical changes outlined the specificity of the interaction between the stimulatory compound and topo II. Based on these observations, we identified two chemical motifs – succinic and glutaric acids – that confer stimulatory activity and conducted a substructure search of these motifs against the Human Metabolome Database. This analysis revealed that the motifs are most strongly represented in TCA cycle intermediates (**Figure 3E and Table S2**). Four TCA cycle intermediates (citrate, isocitrate, succinate, and malate) were identified as stimulatory compounds from our original candidate list (**Table S1**).

### TCA cycle intermediates stimulate topo II activity *in vitro*

Based on our observations, we predicted that other TCA cycle intermediates would have similar effects on topo II activity. All TCA cycle intermediates, with the exception of succinyl-CoA, have succinic and/or glutaric acid substructures (**Figure 3E**). Having ascertained that such a moiety is present in compounds capable of stimulating topo II activity, we proceeded to evaluate the effects of all TCA cycle intermediates on topo II. Consistent with our previous observations, the addition of these intermediates led to as much as a 4- to 12-fold stimulation of decatenation activity and to a 3- to 5-fold stimulation of supercoil relaxation activity at compound concentrations ranging from 25-40 mM (**Figures 4A-J and S3**). Glutamate, an inactive compound (**Figure 3D**), was added as a non-TCA metabolite control to further validate the specificity of the stimulatory effect.

**Figure 4.**
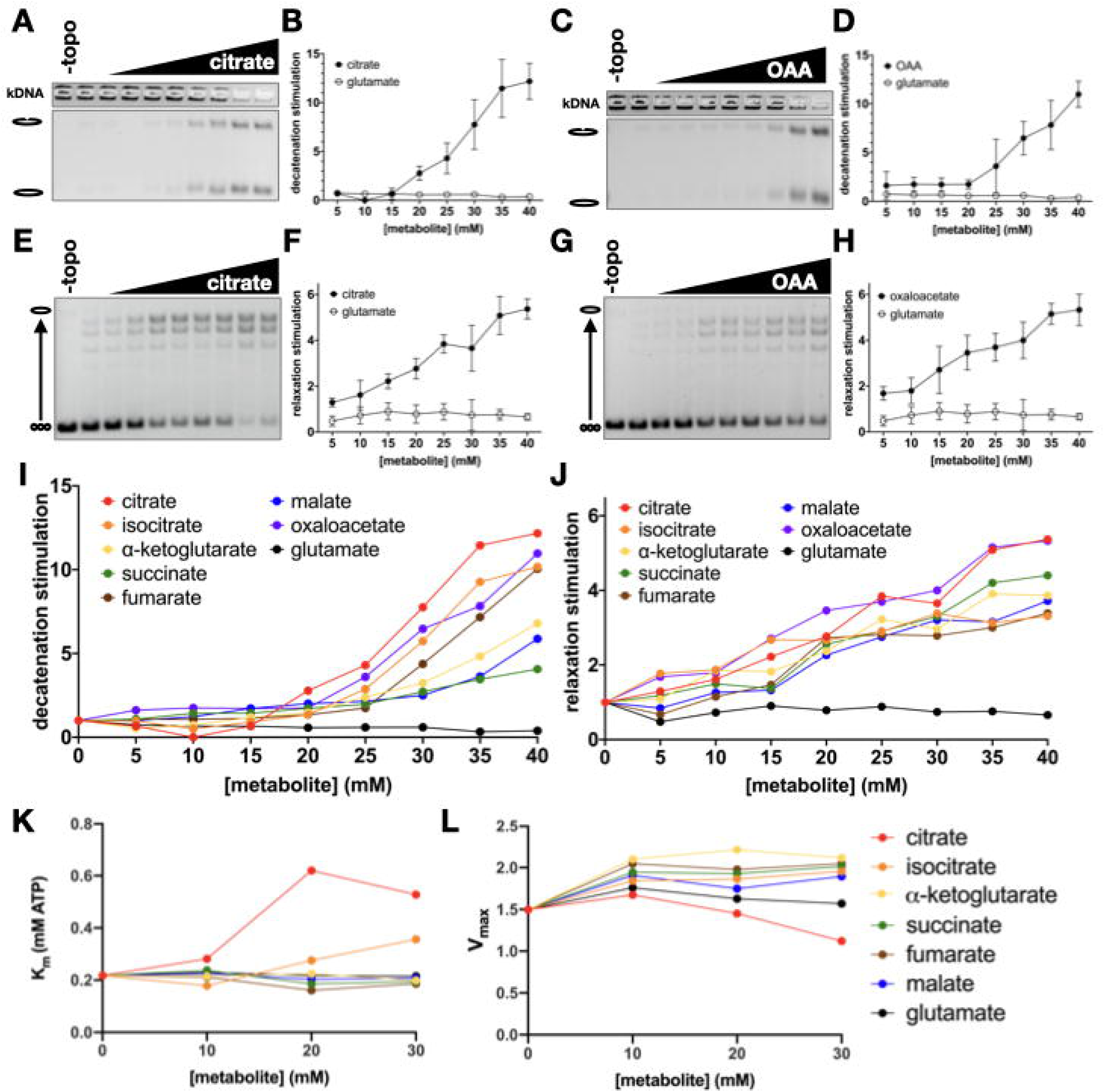
TCA cycle intermediates stimulate strand passage by topo II without significant effects on ATP hydrolysis. (A-H) Stimulation of topo II decatenation activity (A-D) and supercoil relaxation activity (E-H) by citrate and oxaloacetate (OAA). Citrate, OAA, and glutamate (negative control) were titrated from 0 to 40mM in 5mM increments. Graphs (B, D, F, and H) represent mean ± SD of n=3-5 independent experiments. (I-J) Stimulation of *Sc*Top2 decatenation (I) and relaxation (J) activity by TCA metabolites and glutamate (negative control). Stimulation values represent mean of n=3-6 independent experiments. (See also **Figure S3**). (K-L) Effects of TCA metabolites on ATP hydrolysis activity. Experiments were performed at 0, 10, 20, and 30mM concentrations of the metabolites indicated in the legend to the right of the graphs. Km (K) and Vmax (μmol ATP • min^-1^ • nmol topo II^-1^) (L) values were derived from n=3 independent experiments. (See also **Figure S5**).

Of the seven TCA metabolites tested, citrate and isocitrate are distinguished by the presence of a third carboxylic acid group. Citrate is a known Mg^2+^ ion chelator (Yamagami et al., 2018) and Mg^2+^ ions are required for topo II catalytic activity (Osheroff, 1987). To evaluate the potential role of Mg^2+^-ion chelation in the observed stimulation of topo II, we assessed the ability of TCA metabolites to stimulate decatenation activity at varying concentrations of Mg(OAc)_2_ (**Figure S4**). Increasing the concentration of Mg^2+^ alone led to a small but detectable increase in decatenation. However, the addition of 30mM citrate or isocitrate significantly stimulated decatenation beyond the effect of adding Mg^2+^ alone (**Figure S4**). These data indicate that the chelation of free Mg^2+^ ions by citrate or isocitrate does not appreciably contribute to the stimulation of topo II strand passage activity.

We next biochemically characterized the ability of TCA metabolites to modulate the ATPase activity of *Sc*Top2 (**Figures 4K, 4L and S5**). Oxaloacetate was excluded from the ATPase activity experiment because as a substrate for lactate dehydrogenase, it interferes with the coupled reaction. For the evaluated metabolites, we observed that, with the exception of citrate, all led to only a very slight increase in the maximum rate of ATP hydrolysis (< 40%) as compared to the glutamate control. The dicarboxylate intermediates had no significant effect on the K_m_ of enzyme affinity, whereas an increase in K_m_ (∼2-3 fold) was seen with tricarboxylate metabolites (a response that may relate to the chelating activity of these compounds, as Mg^2+^ is a co-factor for the topo II ATPase reaction). Overall, our data indicate that TCA metabolites have relatively little effect on ATP turnover by topo II as compared to the degree of stimulation they exert on the enzyme’s strand passage activity.

### The interaction between topo II and TCA cycle metabolites is specific to eukaryotic type II topoisomerases

Having established a stimulatory interaction between TCA intermediates and budding yeast topo II, we next assessed whether this effect was maintained in human homologs. We tested citrate as a representative tricarboxylic acid metabolite and succinate as representative dicarboxylic acid metabolite against human topo IIα (*Hs*Top2A) and topo IIβ (*Hs*Top2B). Both compounds stimulated DNA decatenation by the human topo IIs (**Figures 5A-H**). *Hs*Top2B decatenation activity was increased by ∼10- to 15-fold by both compounds (**Figures 5A-D)**. Succinate had a similar effect on *Hs*Top2A, whereas stimulation of *Hs*Top2A by citrate was biphasic (**Figures 5E-H**), reaching a peak level of decatenation activity at 30mM citrate that was enhanced ∼20-fold compared to the lowest amount of metabolite tested (5 mM) (**Figure 5F**). Because the TCA cycle is present in prokaryotes as well as eukaryotes, we also tested the activity of citrate and succinate against *E. coli* topo IV (**Figures 5I-L**). Interestingly, both compounds behaved like glutamate and had little to no effect on kDNA decatenation by topo IV. These findings demonstrate that the action of TCA metabolites is specific for eukaryotic type II topoisomerases and evolutionarily conserved from yeast to humans.

**Figure 5.**
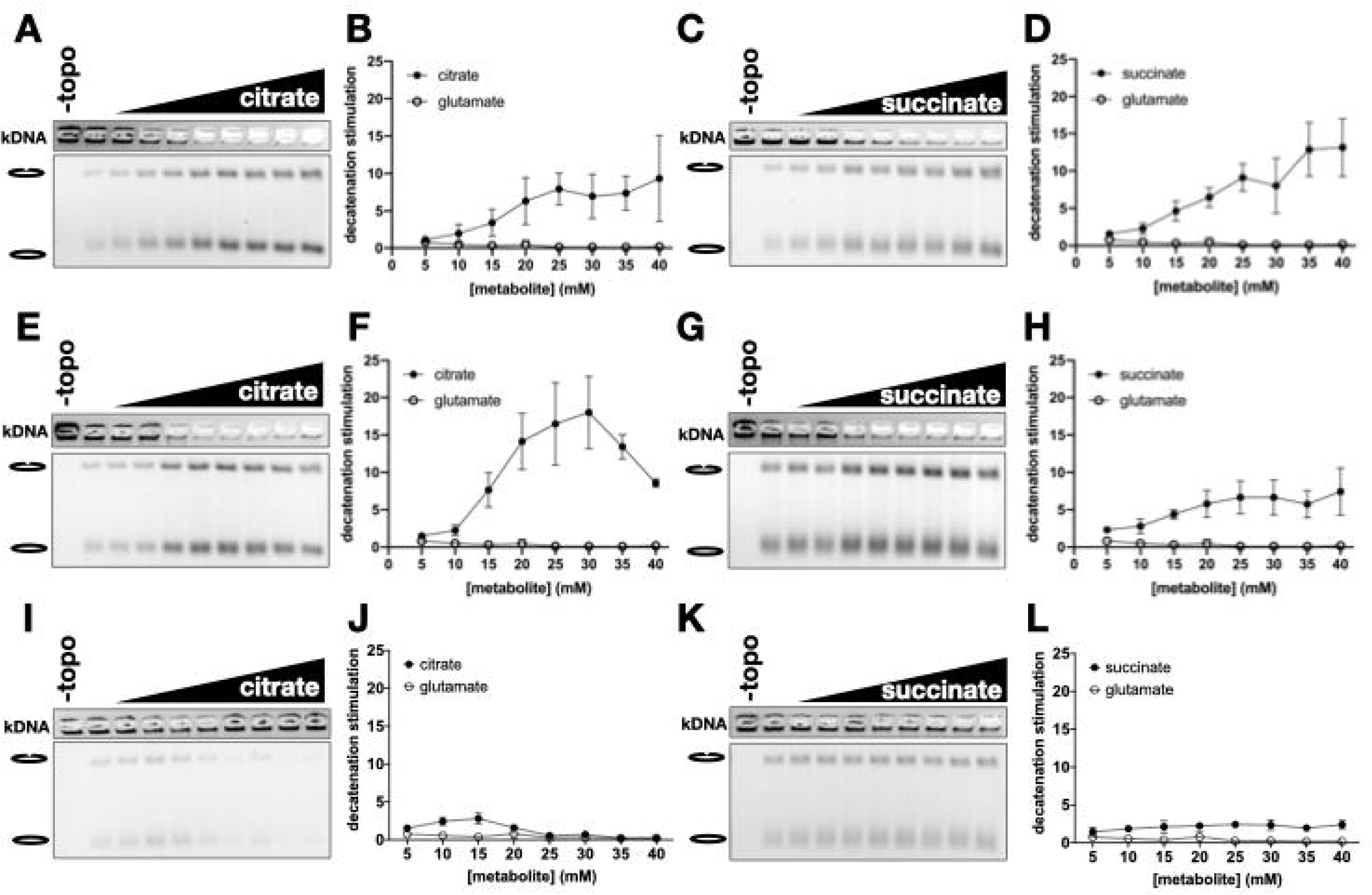
Stimulatory effect of TCA metabolites is eukaryotic specific. Citrate and succinate were titrated from 0 to 40 mM in 5 mM increments and added to decatenation assays with *Hs*Top2B (A-D), *Hs*Top2A (E-H) and *Ec* topo IV (I-L). Graphs represent mean ± SD of n=3-5 independent experiments.

### Altering TCA cycle flux leads to changes in levels of topo II activity *in vivo*

Having established that TCA cycle metabolites exert a stimulatory biochemical effect on topo II, we proceeded to explore the physiological relevance of this activity. Our *in vitro* data predicted that an increase or decrease in the abundance of TCA cycle intermediates should lead to commensurate changes in topo II activity in the cell. To test this hypothesis, we devised two approaches to modulate TCA cycle flux and assess topo II function *in vivo* using drug sensitivity. Experiments were conducted with a drug-efflux deficient (ED) strain of *S. cerevisiae* to improve the dynamic range of the growth assays (Stepanov et al., 2008).

We first assessed the effects of lowered TCA metabolite abundance by deleting *mpc1*, a subunit of the mitochondrial pyruvate carrier (MPC), in the ED background. Transport of pyruvate into the mitochondrial matrix is necessary to replenish TCA cycle intermediates; loss of *mpc1* leads to a significant decrease in the concentration of TCA intermediates (Herzig et al., 2012; Morita et al., 2019). Given our biochemical data, we hypothesized that topo II activity levels should be lower in the *mpc1Δ* strain as compared to the wildtype *MPC1* strain on account of the diminished metabolite levels (**Figure 6A**). To test this idea, we evaluated the sensitivity of the two strains to chemical agents that antagonize topo II. Type II topoisomerase antagonists are broadly classified as either poisons or general catalytic inhibitors (Nitiss, 2009). Poisons stabilize the cleavage complex of topo II bound to substrate DNA, leading the enzyme to form persistent protein-DNA adducts and DNA-strand breaks. As a result, cells that overexpress topo II are hypersensitive to poisoning agents (Nitiss et al., 1992). In contrast, general catalytic inhibitors reduce enzyme activity without directly stimulating DNA breakage. As a consequence of these different modes of action, changes in topo II activity will have opposing effects on the sensitivity of cells to poisons or catalytic inhibitors: increasing topo II activity is expected to lead to hypersensitivity to poisons (due to the elevated formation of toxic cleavage complexes) and resistance to catalytic inhibitors (because stimulation of enzyme activity overcomes general inhibition), whereas decreasing topo II activity should display the converse (**Figure 6A**). Consequently, changes in drug cytotoxicity indicate changes in topo II activity. For our experiments, we chose etoposide as a topo II poison and ICRF-187 as a catalytic inhibitor (Ross et al., 1984; Tanabe et al., 1991). The effects of these drugs on the growth of the *mpc1Δ* strain and the *MPC1* strain were observed by measuring optical density over the course of 35 hours. Interestingly, the *mpc1Δ* strain proved resistant to etoposide and sensitized to ICRF-187 as compared to the control strain, consistent with the idea that decreasing TCA intermediate abundance leads to a corresponding decrease in topo II activity (**Figures 6B, 6C, S6A, and S6B)**.

**Figure 6.**
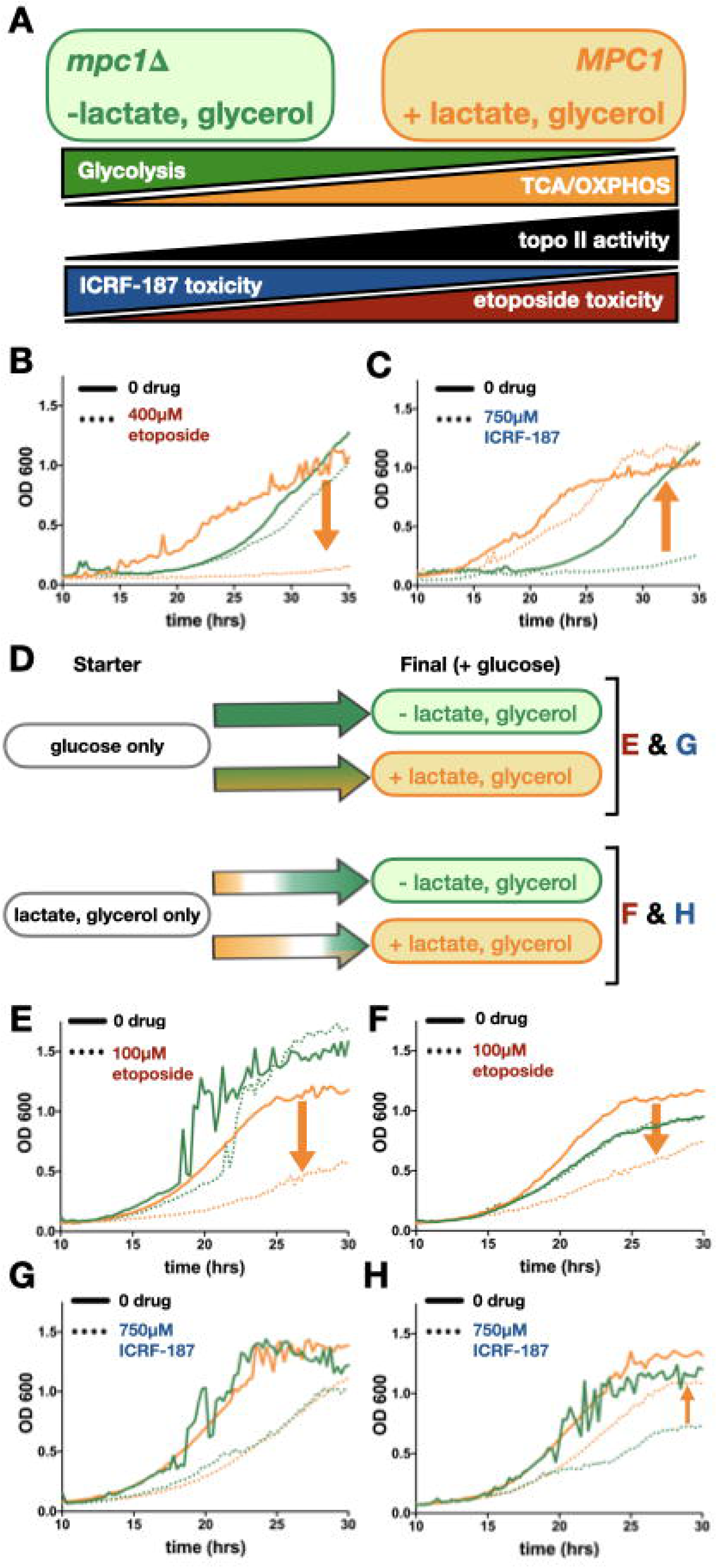
Changes in TCA cycle flux affect sensitivity of yeast to topo II inhibitors. Green indicates conditions in which TCA flux is decreased, while orange indicates conditions in which TCA flux is increased. (A) Predicted correlation between ATP-generating metabolic pathways and topo II activity. Cells generate ATP by glycolysis (green) and oxidative phosphorylation (TCA/OXPHOS, orange). Our model predicts that as TCA metabolism increases, topo II activity will also increase (black). As topo II activity increases, etoposide toxicity (red) increases and ICRF-187 toxicity (blue) decreases; hence, etoposide toxicity should directly correlate with TCA flux and ICRF-187 toxicity should inversely correlate with TCA flux. (B-C) Growth curves of *mpc1* (green) compared to MPC1 (orange) in the presence of etoposide (B) and ICRF-187 (C). Orange arrows show the effect of increasing TCA flux (downward indicates sensitization and upward indicates rescue). (See also **Figure S6A-B**) (D) Schematic depiction of the effects of nutrient changes on flux through glycolysis and TCA cycle. Arrows are colored to show shifts in metabolism over time. Cultures initially grown in glucose conditions (E, G) will continue to have high levels of glycolysis, but the addition of lactate and glycerol will cause an increase in TCA flux. Cultures initially grown in media with only lactate and glycerol (F, H) will generate ATP solely through TCA/OXPHOS. Introduction of glucose will cause a diauxic shift towards glycolysis, but if lactate and glycerol are maintained in the media, this shift will be delayed and TCA flux will stay elevated. Letters to the right of (D) indicate the nutrient conditions of the growth curves shown in (E-H). (E-H) Green lines show growth in glucose only media and orange lines show growth in media with glucose, lactate, and glycerol. Cultures were inoculated from starters grown in glucose only media (E and G) or lactate and glycerol only media (F and H). Orange arrows show the effect of increasing TCA flux on cytotoxicity of etoposide (E and F) and ICRF-187 (G and H) (downward indicates sensitization and upward indicates rescue). (See also **Figure S6F-I**).

To test the effect of increasing TCA metabolite abundance, we developed a strategy to enhance TCA cycle flux by adding non-fermentable carbon sources to the growth media. As described by the Crabtree effect (Crabtree, 1928), yeast will opt to generate ATP through glycolysis rather than oxidative phosphorylation under high glucose conditions, regardless of oxygen availability. However, when fermentable carbon sources are limited, yeast will generate ATP by oxidative phosphorylation (OXPHOS), which is coupled to the TCA cycle. To induce different metabolic states, we cultured yeast in minimal media either with glucose only, or with glucose supplemented with lactate and glycerol (LG). Lactate and glycerol are non-fermentable and must be metabolized through the TCA cycle to generate ATP. By LC-MS analysis, we were able to qualitatively compare the levels of five TCA metabolites in yeast grown glucose media with and without LG. We detected significant increases in succinate and malate in the presence of LG and no significant changes in the other metabolites (**Figure S6C).** To further check that the addition of LG increases TCA flux, we compared the effect of UK-5099, a chemical inhibitor of the MPC that inhibits pyruvate entry into the TCA cycle, on colony growth in the two different nutrient environments. Yeast cells proved more resistant to UK-5099 in media containing LG as compared to the glucose-only condition, confirming that the addition of non-fermentable carbon sources increases TCA flux to counteract the effects of UK-5099 (**Figure S6D**).

We initially examined the effects of adding LG on the sensitivity of yeast cells to anti-topo II agents using a spot-growth assay. The addition of LG sensitized yeast to etoposide as expected (**Figure S6E**) but we were unable to reach cytotoxic concentrations of ICRF-187 necessary to see rescue. We therefore designed a liquid media growth assay that incorporates a diauxic shift to assess the impact of TCA metabolism on cellular sensitivity to topo II antagonists (**Figure 6D)**. Starter cultures were first grown to saturation in media containing either glucose alone, or in LG alone to allow the yeast to adjust to either a glycolysis-dominant or an OXPHOS-dominant expression profile. These cells were then shifted into glucose-containing media with and without LG to monitor growth in the presence of anti-topo II drugs. Because glucose is the preferred carbon source, yeast from the glucose-only starter condition will primarily rely on glycolysis and show only a slight increase in TCA metabolism in the presence of LG (**Figure 6D**). By contrast, yeast grown with only non-fermentable carbon sources will undergo a diauxic shift as glucose is introduced. If LG are present, this diauxic shift will be delayed and the concentration of TCA intermediates will be elevated as the cells attune to metabolizing non-glycolytic carbon substrates (**Figure 6D**). In agreement with the spot test, the addition of LG sensitized cells to etoposide (**Figures 6E, 6F, S6F, and S6G**). The addition of LG did not rescue cells that were acclimated to glucose only nutrient conditions from ICRF-187 (**Figures 6G and S6H**). However, the addition of LG did partially rescue yeast from ICRF-187 toxicity when the starter culture was grown without glucose (**Figures 6H and S6I**). Collectively, these results show that cellular topo II activity correlates with changes in TCA cycle flux.

To confirm that the changes in sensitivity to topo II-targeted drugs were due to catalytic stimulation and not a result of differences in protein expression levels, we generated two yeast strains with a 3xHA-tag at either the N-terminal or C-terminal end of endogenous topo II. These strains were then cultured in glucose-only media or glucose media supplemented with LG to compare topo II expression levels under these different growth conditions (**Figure S7**). Cells were harvested in mid-log phase to capture topo II expression in actively dividing cells and assessed for topo II levels by Western blot. These data show that the change in relative protein abundance across the different growth conditions was insignificant, supporting the idea that the differences in topo II activity observed from the drug screen derive from biochemical stimulation of the enzyme and not due to alterations in protein concentration.

## DISCUSSION

### Regulatory interactions between endogenous metabolites and DNA machinery

In the present work, we show that DNA topoisomerase II (topo II) is subject to regulatory control by small-molecule intermediates of the TCA cycle. Short-chain (2-and 3-carbon spaced) di- and tricarboxylic acids are found to stimulate both DNA supercoil removal and DNA decatenation by topo II *in vitro* (**Figure 4**). Interestingly, TCA cycle intermediates stimulate strand passage without increasing ATP turnover rate (**Figures 4 and S5**), indicating that these agents increase enzyme efficiency. Human topo IIs are also stimulated by TCA intermediates, whereas homologous bacterial type II topoisomerases are not, suggesting that topo II evolved a response to metabolic control to meet a specific regulatory need in eukaryotes (**Figure 5**).

TCA cycle intermediates are some of the most abundant metabolites in the cell, with some intermediates such as citrate, succinate, and malate each reaching concentrations in the 1-5 mM range (Park et al., 2016; Wittmann et al., 2005). Given that seven out of eight TCA cycle intermediates stimulate to topo II activity, a global shift in the collective concentration of this metabolite pool would be expected to exert a functional effect. *In vitro*, supercoil relaxation was stimulated ∼5-fold and DNA decatenation ∼12-fold stimulation at the higher metabolite concentrations tested (≥ 20 mM) (**Figure 4**). However, up to 3-fold increases in activity were still evident even at lower, more physiological levels (between 5-15mM). Many biological systems are maintained at homeostatic setpoints where modest (sub-twofold) biochemical changes can have profound physiological consequences. For example, a < 2-fold increase in *K_m_* and < 20% decrease in *k_cat_* of the tetracycline resistance protein, TetX2, is sufficient to impart robust resistance to this antibiotic in bacteria (Walkiewicz et al., 2012). Similarly, a ∼15% decrease in the expression of mismatch repair protein MSH2 is reported to confer chemoresistance to the chemotherapeutic, temozolomide, in glioblastoma cells (McFaline-Figueroa et al., 2015). Regulatory processes can also be highly sensitive to minor changes in protein activity. A < 10% difference in the concentration of Bicoid, a morphogen, determines cell fate during *D. melanogaster* embryonic development through transcriptional regulation (Driever and Nüsslein-Volhard, 1988; Fradin, 2017). Based on these examples, it is not unreasonable that even a two-fold change in topo II activity resulting from differing TCA cycle intermediate concentrations would have physiologic consequences. Consistent with this idea, we show that TCA cycle status indeed directly impacts topo II function *in vivo*, with changes in TCA flux altering the relative sensitivity of budding yeast to two different classes of topo II-targeted drugs (**Figure 6**).

Why link topo II activity to metabolic status? One possibility is that it may provide a means to coordinate enzyme function with the cell cycle. Cell cycle-dependent variation in topo II concentration is already known to occur for topo IIα in eukaryotes that express two isoforms of the enzyme (Heck et al., 1988; Kimura et al., 1994; Woessner et al., 1991). Too little or too much topo II activity throughout the cell cycle can impair chromosome condensation and sister chromatid segregation, potentially leading to cell-cycle arrest (Andrews et al., 2006; Cuvier and Hirano, 2003; Downes et al., 1994; Giménez-Abián et al., 1995; Samejima et al., 2012; Uemura et al., 1987). Metabolic sensing by topo II may provide a means for fine-tuning activity, as metabolism is highly responsive to environmental changes (Brauer et al., 2008; Lee and Finkel, 2013; Wellen and Thompson, 2010; Zhu and Thompson, 2019).

The cell cycle is a controlled sequence of events resulting in the duplication of cellular biomass and faithful segregation of genetic material (Elliott and McLaughlin, 1978; Goranov et al., 2009; Johnston et al., 1977; Jorgensen et al., 2002; Kalucka et al., 2015). This process is not only energetically demanding but also requires the activation of anabolic pathways to generate macromolecular building blocks necessary to accommodate increased transcription, translation, and DNA replication. As such, metabolism has been observed to fluctuate in accordance with cell-cycle progression, and cell-cycle regulators such as cyclin/CDK complexes and the APC-C/Cdh1 have been shown to coordinate metabolic shifts (Almeida et al., 2010; Buchakjian and Kornbluth, 2010; Colombo et al., 2010; Liu et al., 2020; Salazar-Roa and Malumbres, 2017; Tudzarova et al., 2011; Wang et al., 2014). Proliferating cells strongly up-regulate glycolysis in G1 to rapidly generate ATP and carbon substrates for anabolic pathways (Buchakjian and Kornbluth, 2010; Diaz-Moralli et al., 2013; Pavlova and Thompson, 2016), but as cells enter S phase, glycolysis is down-regulated and metabolism shifts towards the pentose phosphate pathway to support DNA replication (Diaz-Moralli et al., 2013; Da Veiga Moreira et al., 2015). As transcription ramps down upon chromosome condensation (Johnson and Holland, 1965; Littau et al., 1964; Prescott and Bender, 1962; Taylor, 1960), the cell shifts towards the TCA cycle and OXPHOS as a more efficient source of ATP to fuel mitotic progression (Wang et al., 2014). The shift in energy metabolism from glycolysis towards TCA cycle/OXPHOS at mitotic entry mirrors the upregulated expression pattern of vertebrate topo IIα that occurs at the same cell cycle transition point. Based on these observations we hypothesize that TCA cycle fluctuations coordinate an increase of topo II activity at the G2-to-mitosis transition, likely to help ensure that chromosomes are fully decatenated. This control mechanism may be particularly important in lower-order eukaryotes that do not display cell cycle-dependent expression of topo II and could have been preserved in vertebrates that are capable of upregulating topo IIα expression as means of contending with the large chromosomal content.

### Cancer specific metabolic profiles may impact efficacy of topo II-targeted chemotherapeutics

Cancer cells have distinguishing metabolic features, one of the first of which was observed by Otto Warburg (Warburg, 1925). The Warburg Effect, characterized by increased aerobic glycolysis and lactic acid fermentation in cancer cells, was initially thought to indicate a shift away from TCA cycle metabolism and OXPHOS to favor the rapid production of ATP through glycolysis; however, growing evidence shows that increased rates of glycolysis serve to provide anabolic intermediates for biosynthetic pathways, and that the TCA cycle and OXPHOS both remain active in cancer cells (Locasale et al., 2011; Pavlova and Thompson, 2016). Diverting pyruvate away from the TCA cycle through lactic acid fermentation replenishes NAD+ to maintain high rates of glycolysis (Luengo et al., 2020; Pavlova and Thompson, 2016). TCA cycle metabolites are key anabolic precursors and citrate, in particular, is critical to support elevated levels of lipid biosynthesis in cancer cells (Li and Cheng, 2014; Menendez and Lupu, 2007). Recent studies have also found that the impairment of TCA cycle metabolism and OXPHOS can decrease tumorigenic and metastatic potential (Cai et al., 2020; Cavalli et al., 1997; Lebleu et al., 2014; Morais et al., 1994; Tan et al., 2015). These findings demonstrate that cancer cells not only maintain active TCA cycle metabolism but are dependent on it.

Different cancer cell types have been shown to have distinct metabolic dependencies that may influence their response to various treatment modalities (Diaz-Ruiz et al., 2011; Guppy et al., 2002; Martin et al., 1998; Pasdois et al., 2003). With respect to the TCA cycle, mutations in several key TCA cycle enzymes have been observed in specific cancers. Isocitrate dehydrogenase (*IDH1/2*) mutations are frequently found in gliomas, cholangiocarcinomas, and a subset of acute myeloid leukemias (Balss et al., 2008; Borger et al., 2012; Yan et al., 2009; Yang et al., 2012). In such cases, a single amino acid substitution at Arg132 causes IDH to generate 2-hydroxyglutarate (2HG) rather than its normal product, α-ketoglutarate (Dang et al., 2009; Ward et al., 2010; Yang et al., 2012). Interestingly, 2HG was identified as a topo II-stimulating metabolite in our studies (**Table S1**), suggesting that cancers that accumulate 2HG may have elevated levels of topo II activity. Inactivating mutations in succinate dehydrogenase (SDH) and fumarate hydratase (FH) have also been shown to cause an accumulation of succinate and fumarate in paragangliomas and pheochromocytomas, as well as in some gastrointestinal and renal cell cancers (Astuti et al., 2001; Baysal et al., 2000; Janeway et al., 2011; King et al., 2006; Letouzé et al., 2013; Selak et al., 2005; Tomlinson et al., 2002). The accumulation of oncometabolites (e.g., 2HG, succinate, and fumarate) are thought to affect cell fate determination pathways by disrupting chromatin modification status and DNA repair processes (Baksh and Finley, 2020; Intlekofer and Finley, 2019; Sulkowski et al., 2020; Xiao et al., 2012). Our demonstration that elevated levels of TCA intermediates sensitize yeast cells to topo poisons (**Figure 7**) suggests that some topo II-targeted therapies may be more effective and selective against cancers with that accumulate oncometabolites. Future efforts will be needed to test this supposition.

**Figure 7.**
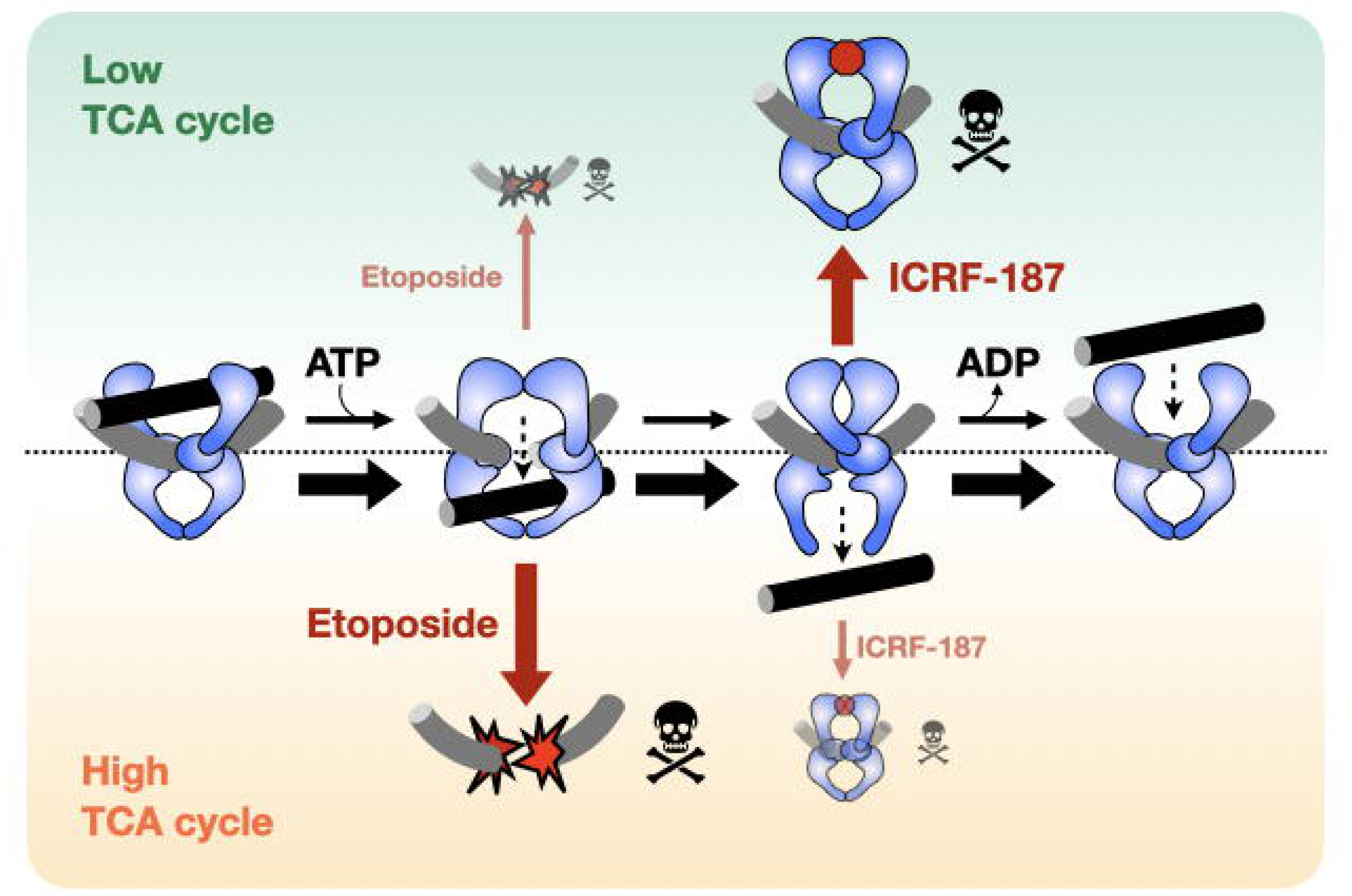
Schematic depicting how sensitivity to topo II targeting drugs is influenced by metabolic state. When TCA cycle flux is low (green) topo II strand passage activity is not stimulated. General catalytic inhibitors (e.g., ICRF-187) that work by decreasing topo II activity will be more effective in this state, as compared to a high TCA cycle flux state (orange). By contrast, topo II poisons (e.g., etoposide) are more toxic when topo II activity is stimulated in the high TCA cycle flux state because elevated topo II strand passage activity leads to increased formation of cytotoxic DNA double strand breaks.

### Concluding remarks

The present work reports on the ability of native small-molecule metabolites to directly activate eukaryotic topo II *in vitro* and *in vivo*. Although there exist many different classes of topo II antagonists, this is to our knowledge the first report of a small molecule capable of stimulating this enzyme. Polyamines, such as spermine and spermidine, have been shown to enhance topo II supercoil relaxation activity by condensing DNA but these compounds do not directly bind topo II (Pommier et al., 1989; Srivenugopal et al., 1987). Because polyamines act on the DNA substrate and not the enzyme, they also affect the activity of type I topoisomerases and bacterial type II topoisomerases (Srivenugopal and Morris, 1985; Srivenugopal et al., 1987). By contrast, the specificity of TCA metabolites for stimulating eukaryotic topo II indicates that the enhancement of strand passage activity is due to a direct interaction between the enzyme and the metabolites (**Figure 5**). These findings also represent the first example of an allosteric interaction between a non-nucleotide metabolite and an enzyme involved in controlling a ‘central dogma’ process. Small molecule ‘alarmones,’ such as (p)ppGpp and cGAMP, do exist that can regulate translation, transcription, and DNA replication (Srivatsan and Wang, 2008; Sun et al., 2013); however, these agents are nucleotide analogs and are produced in response to stress, as opposed to the normal operations of the cell. Metabolites such as α-ketoglutarate, succinate, fumarate, and 2HG can act as cofactors or competitive inhibitors for chromatin modifying enzymes (Baksh and Finley, 2020; Intlekofer and Finley, 2019), but prior to this study, none were known the allosterically regulate an enzyme involved in nucleic acid transactions. The discovery of a regulatory topo II-TCA cycle interaction opens up the possibility that other such interactions between nuclear enzymes and endogenous small molecules may exist for the purposes of modulating events such as transcription and/or DNA replication. Our approach for isolating metabolite-enzyme interactions is readily transferable to any system where enzyme activity can be monitored by a biochemical assay. We expect that future applications of the approach may be useful for revealing other regulatory connections between cellular metabolism and DNA-dependent machineries.

With regard to anti-topo II agents as cancer therapeutics, the off-target poisoning of topo IIβ appears to be at least one major source of negative side effects, including cardiotoxicity and development of secondary malignancies (Azarova et al., 2007; Felix, 1998; Turcotte et al., 2018; Yi et al., 2007). Because of the high degree of similarity between the two human isoforms of topo II, designing strategies to increase the specificity of topo II inhibitors is challenging. Our data show that there is a difference in the biochemical response of topo IIα and topo IIβ to di- and tricarboxylic acids (**Figure 5**); defining the biochemical underpinnings of these differences may provide new insights toward increasing the specificity of topo II-targeted drugs. To this end, we note that some fractions of our metabolic extracts also showed inhibitory activity against topo II, indicating that there exist other biologically relevant metabolite interactions that have yet to be uncovered. The identification of these agents could serve as a starting point for such investigations.

In closing, the relationship between TCA cycle metabolism and topo II activity gives rise to many new questions about how topo II drugs interact with cancerous and noncancerous cells. A deeper understanding of the metabolic regulation of topo II activity may be useful for improving for patient selection strategies and topo II-targeted chemotherapies. Future studies expanding into mammalian systems will clarify the conserved link between cellular metabolism and topo II regulation with the ultimate goal of improving the safety and efficacy of patient-specific cancer treatments.

## Acknowledgments

We thank B. Cormack and members of C. Wolberger’s lab for access to and guidance on the usage of equipment required for metabolite purification and liquid culture yeast growth assays. We also thank M. Bjornsti for sharing the drug efflux deficient yeast strain and G. Hauk for providing purified *E. coli* topo IV. We are grateful to J. Stivers, B. Cormack, and T. Lee for useful advice in carrying out this effort, as well as M. Wolfgang and S. Yegnasubramanian for critical reading of the manuscript. These studies were supported by the NIH (T32-GM007445 and F31-CA224896 to JL, R01-CA168653 to YSL, R01-GM103853 to NNB, and R01-CA077373 to JMB).

## Author Contributions

JHL conducted *in vitro* assays, fractionated yeast metabolites, and performed yeast cell growth assays. EPM conducted LC-MS/MS analysis of metabolite samples. Data analysis was performed by JHL and EPM. YSL contributed to the development of the yeast metabolite extraction method. JHL, EPM, NNB, and JMB contributed to conceptual planning, experimental design, and data interpretation. JHL, EPM, and JMB prepared the manuscript. All authors critically reviewed the manuscript and approved the final version.

## Supplemental Figure Legends

**Figure S1.**
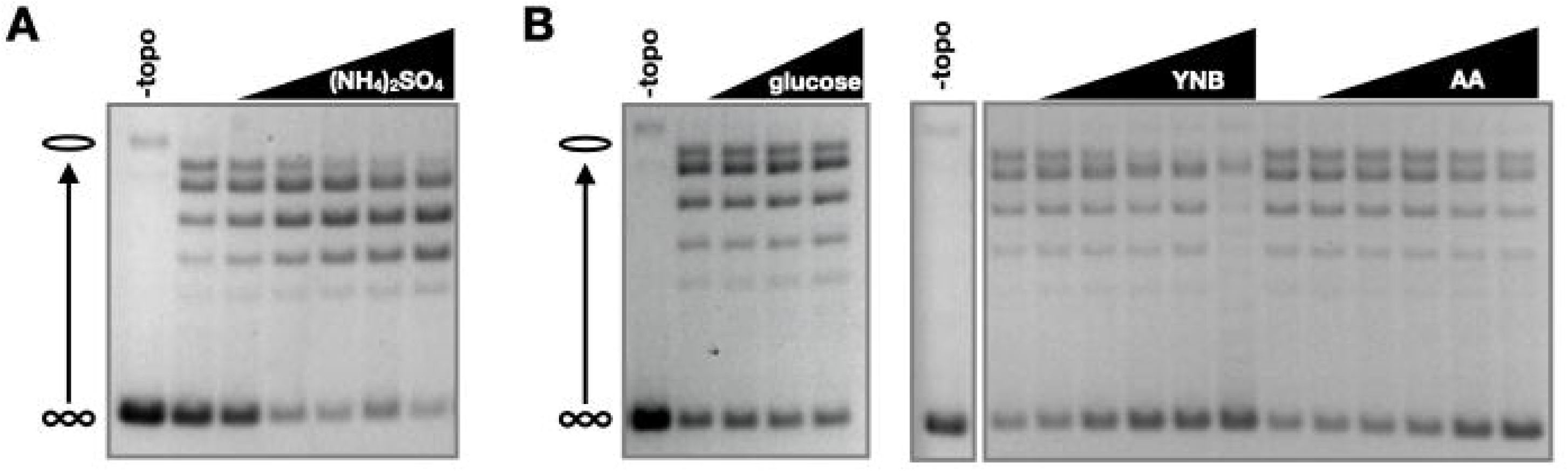
Effects of synthetic media components on topo II activity. (A) Schematic for acid/base butanol liquid-liquid extraction of crude metabolite extracts. ‘Aq’ indicates the aqueous fraction, and ‘Org’ indicates the organic fraction. (B) Supercoil relaxation assay with fractions from acid/base butanol extraction as indicated by colors. Solid material left after solvent removal from each fraction was solubilized in equal volumes of 12.5% DMSO and titrated into relaxation assays in 2-fold dilution steps. (C-D) Activity of crude metabolites after enzymatic treatment by Antarctic phosphatase (C) or snake venom phosphodiesterase (D). Lyophilized material from treated and untreated samples were solubilized in equal volumes of 12.5% DMSO and titrated into relaxation assays in 2-fold dilution steps.

**Figure S2.**
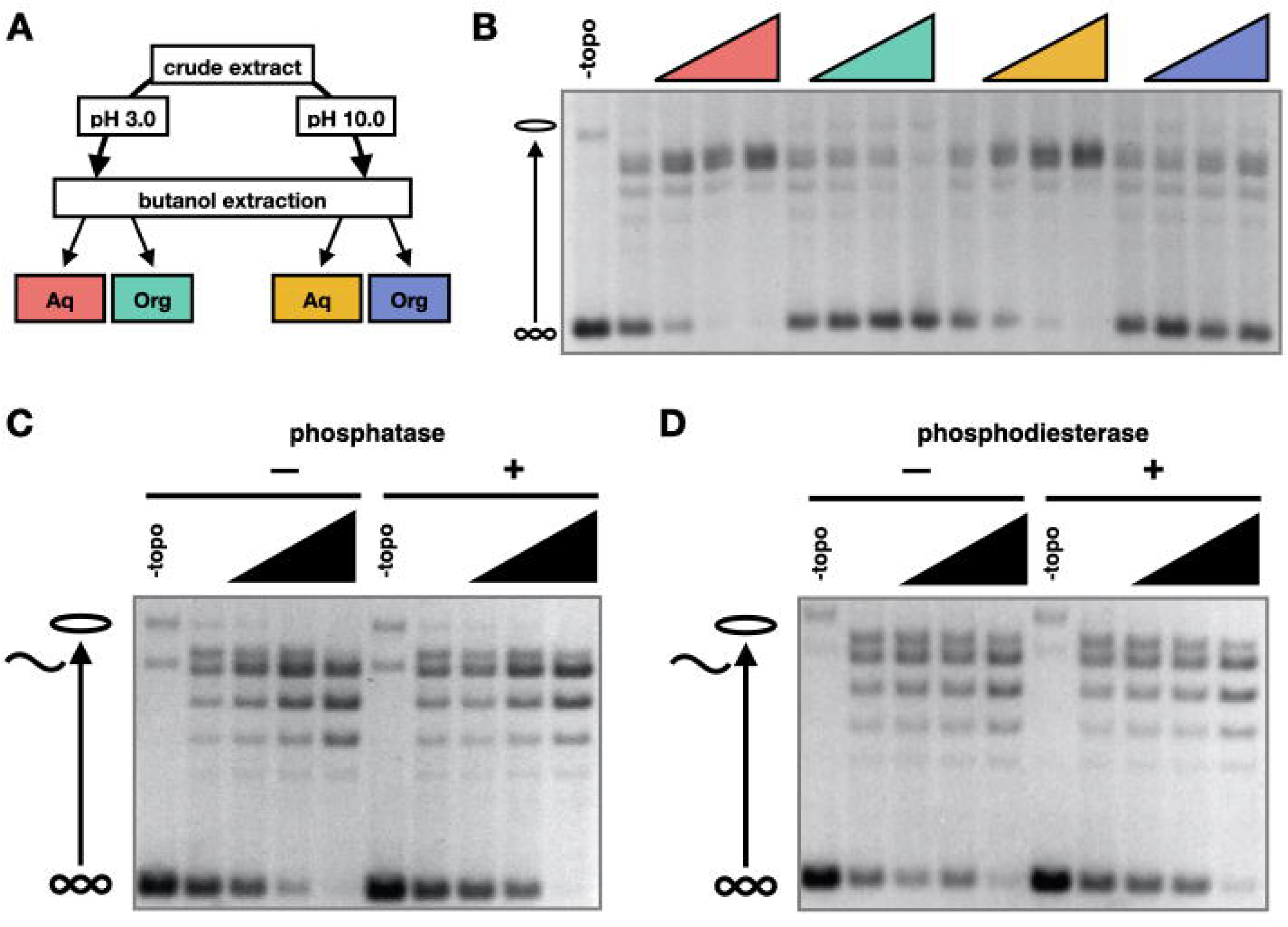
Chemical properties of stimulatory metabolites. (A) Schematic for acid/base butanol liquid-liquid extraction of crude metabolite extracts. ‘Aq’ indicates the aqueous fraction, and ‘Org’ indicates the organic fraction. (B) Supercoil relaxation assay with fractions from acid/base butanol extraction as indicated by colors. Solid material left after solvent removal from each fraction was solubilized in equal volumes of 12.5% DMSO and titrated into relaxation assays in 2-fold dilution steps. (C-D) Activity of crude metabolites after enzymatic treatment by Antarctic phosphatase (C) or snake venom phosphodiesterase (D). Lyophilized material from treated and untreated samples were solubilized in equal volumes of 12.5% DMSO and titrated into relaxation assays in 2-fold dilution steps.

**Figure S3.**
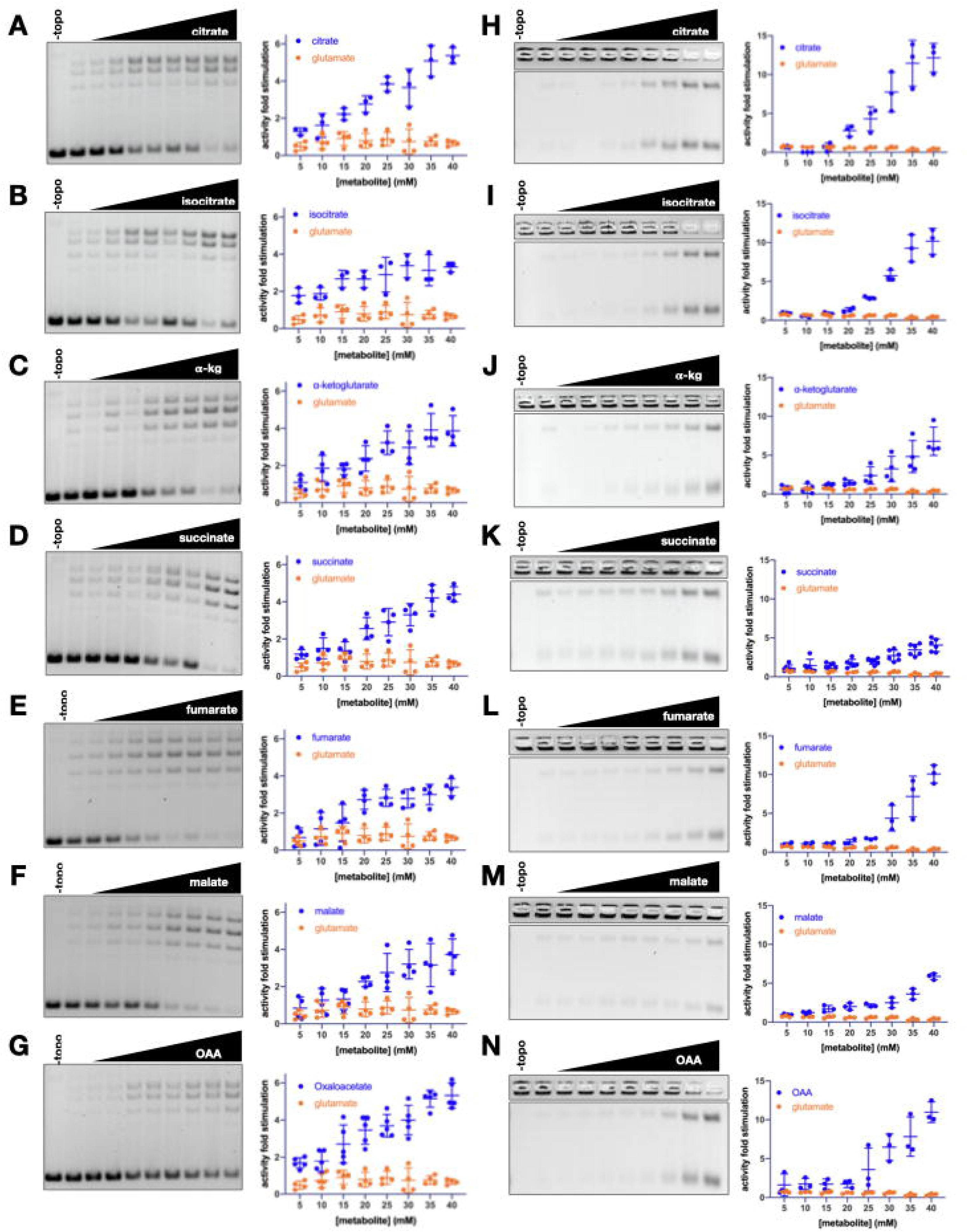
Stimulation of *Sc*Top2 supercoil relaxation activity (A-G) and decatenation activity (H-N) by TCA cycle intermediates (blue) and glutamate (negative control, orange). Graphs show mean ± SD of n=3-6 independent experiments.

**Figure S4.**
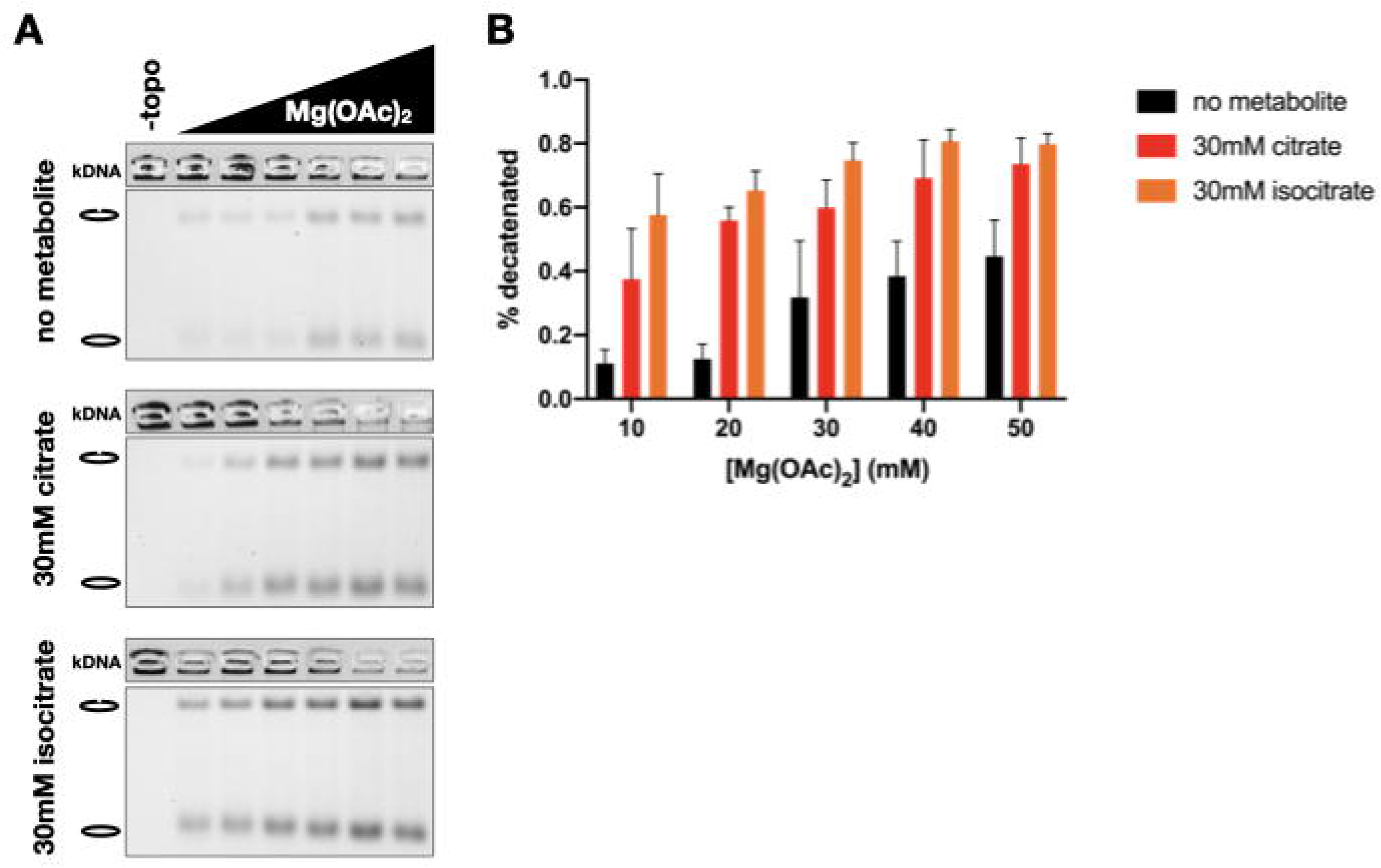
Chelation of Mg^2+^ ions by tricarboxylate TCA cycle intermediates does not affect stimulatory topo II-metabolite interaction. (A) Representative gels of decatenation assays performed at different concentrations of Mg(OAc)_2_ with no added metabolite, 30mM citrate, and 30mM isocitrate. Mg(OAc)_2_ was titrated from 10 to 50 mM in 10 mM steps. (B) Graphs show mean ± SD of n=3 independent experiments to compare the fraction decatenated under each reaction condition.

**Figure S5.**
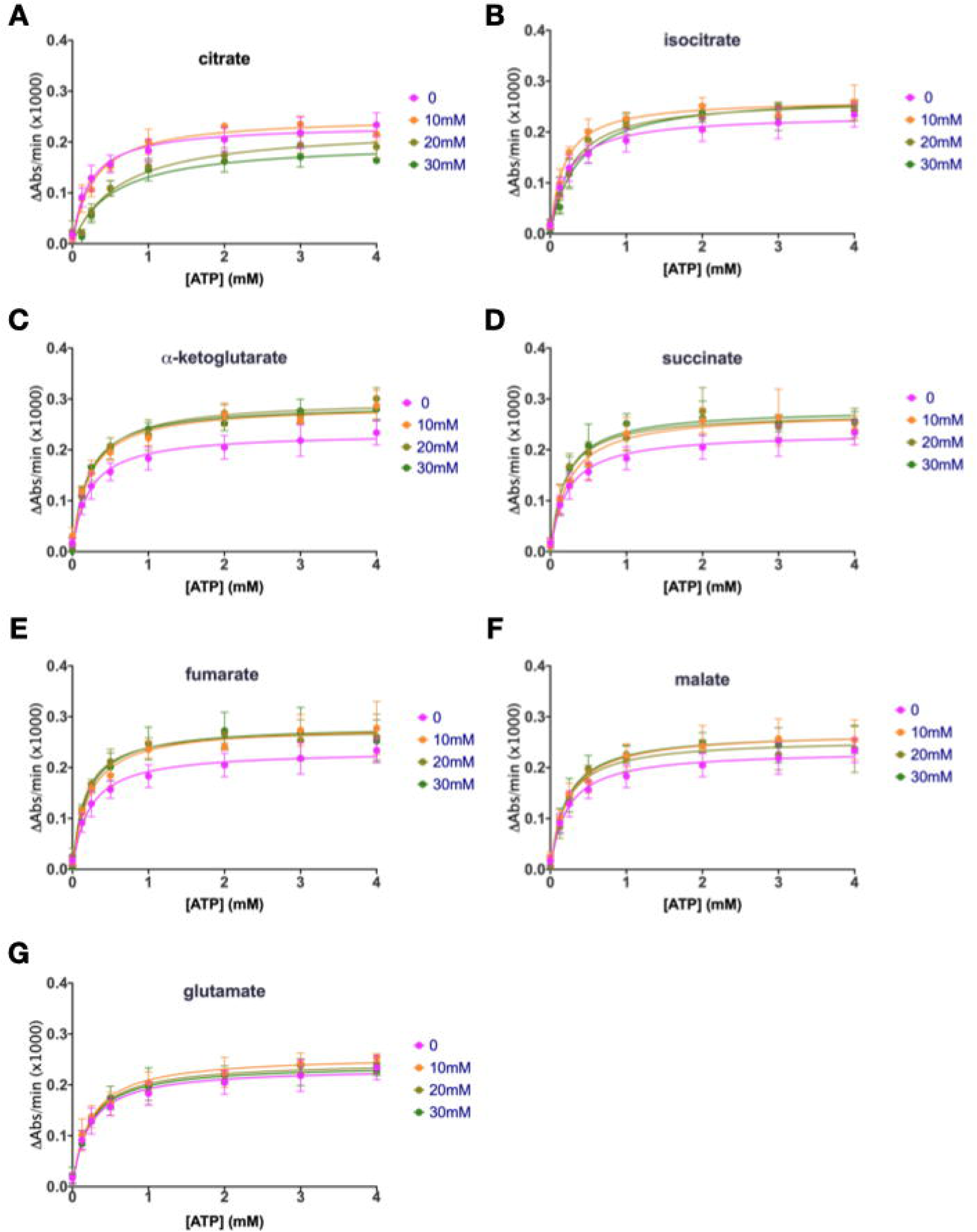
Michaelis-Menten curves of ATP hydrolysis by *Sc*Top2 in the presence of TCA cycle intermediates and glutamate (negative control). Rates of ATP hydrolysis were measured in a coupled assay by monitoring NADH consumption as a function of light absorbance at 340 nm wavelength (y-axis). Graphs represent mean ± SD of n=3 independent experiments.

**Figure S6.**
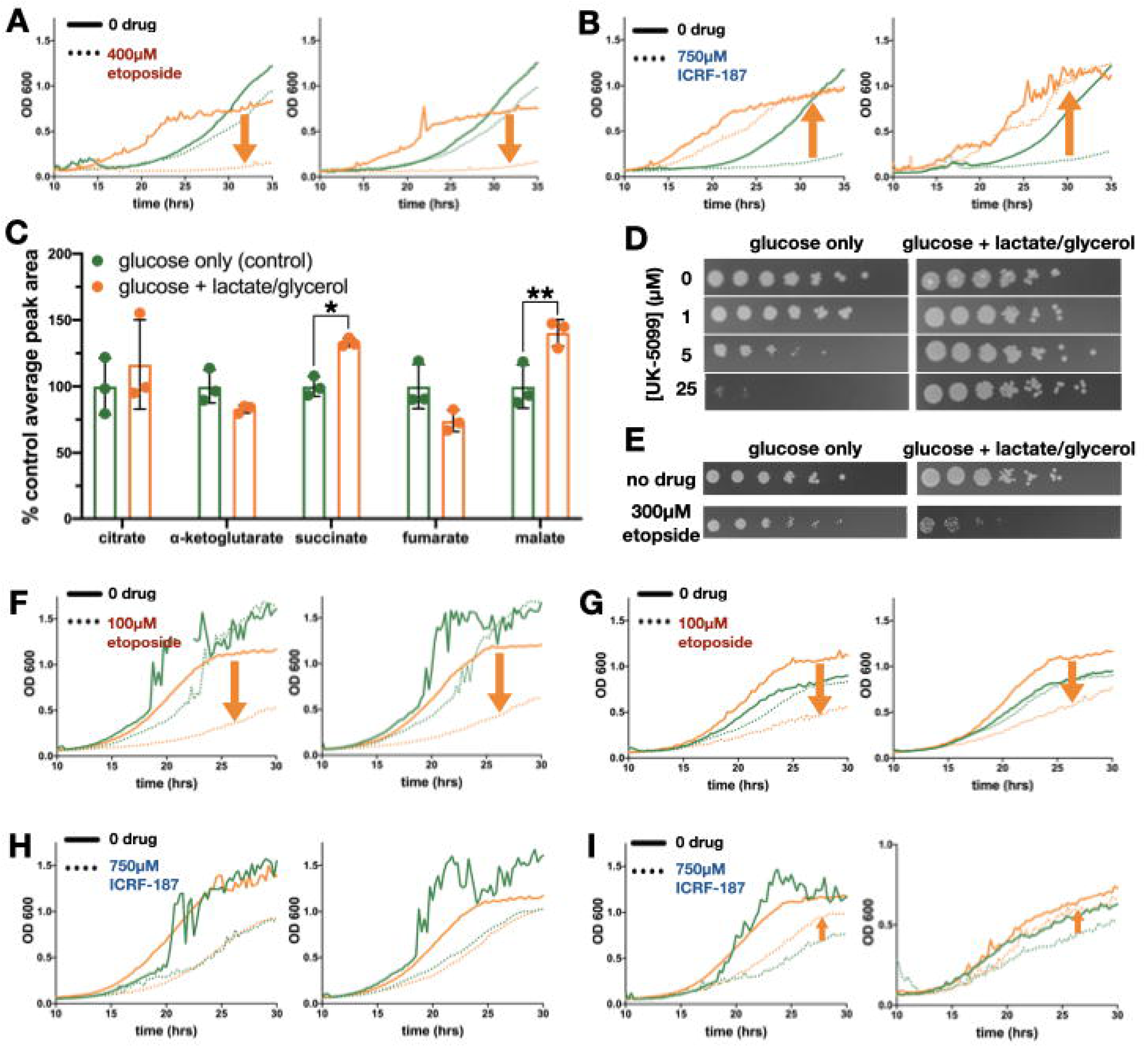
Changes in TCA cycle flux affect sensitivity of yeast to topo II inhibitors - extended data. (A-B) Growth curves of MPC1 (orange) and *mpc1* (green) with etoposide (A) and ICRF-187 (B). (C) Changes in TCA cycle metabolite abundance upon addition of lactate and glycerol to media. Peak areas were normalized to the average of the control condition (glucose only). Bar graphs and error bars indicate mean ± SD of n=3. An unpaired, two-tailed t-test was used to analyze significance of changes peak area. One asterisk (*) indicates p < 0.05, and two asterisks (**) indicates p < 0.01. (D-E) Serially diluted cultures of yeast were spotted on glucose containing agar plates M UK-5099 (D) and 300 μM etoposide (E) was observed after 2-3 days. (F-H) Growth curves of yeast in glucose-only media (green) or glucose media supplemented with lactate and glycerol (orange). Cultures were inoculated from starters grown in glucose only media (F and H) or lactate and glycerol only media (G and H). Orange arrows show the effect of increasing TCA flux on cytotoxicity of etoposide (F and G) and ICRF-187 (H and I) (downward indicates sensitization and upward indicates rescue).

**Figure S7.**
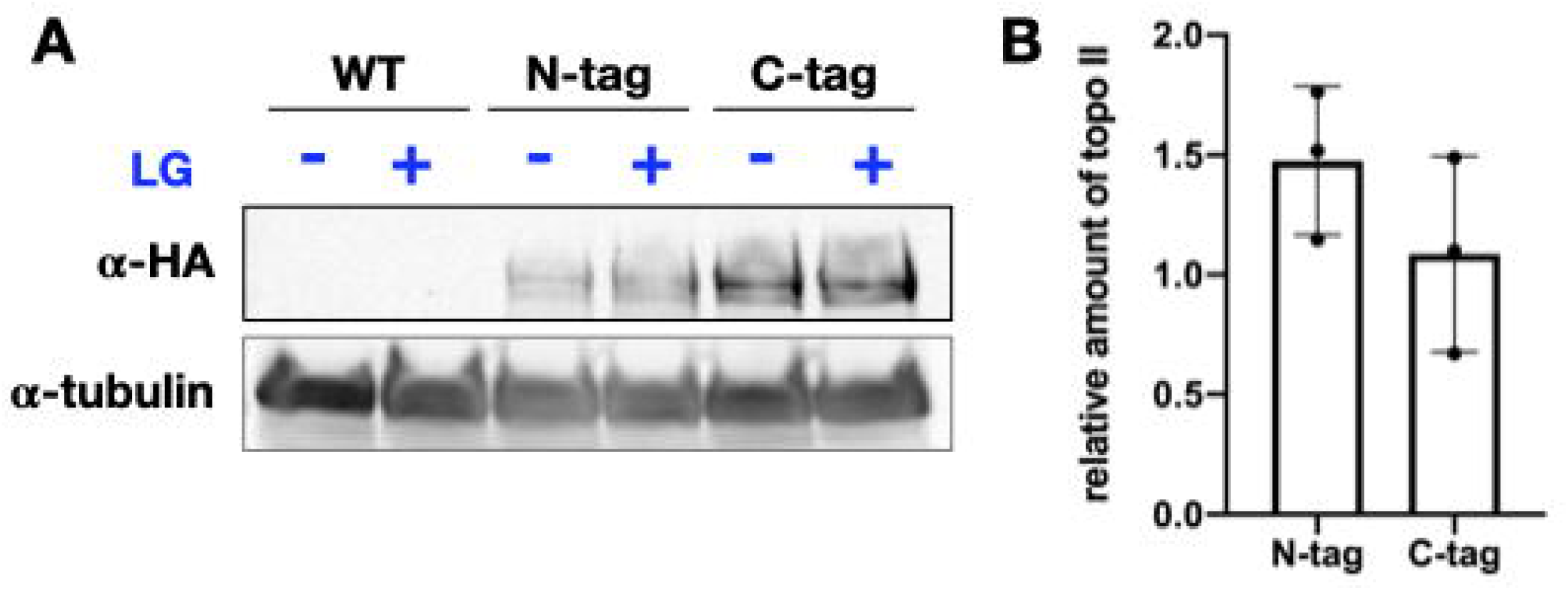
*Sc*Top2 expression levels in different nutrient conditions. (A) Representative western blot of 3xHA-tagged endogenous topo II in an N-terminally tagged (N-tag) or a C-terminally tagged (C-tag) topo II strain from cells grown in media with and without lactate and glycerol (LG). The wildtype (WT) yeast strain (untagged topo II) is shown as a negative control. (B) Quantification of topo II expression levels in the N-tag and C-tag strains. Intensities of the topo II bands were normalized to the corresponding tubulin band. Relative amounts of topo II were calculated by dividing the normalized topo II intensity in the +L/G condition to that of the - L/G condition. Data are represented as mean ± SD of n=3 independent experiments.

Table S1. **Relative abundance and activities of metabolites identified in LC-MS/MS analysis of inactive, stimulatory, and inhibitory fractions of yeast metabolite extracts.**

Table S2. **Results of substructure search of succinic and glutaric acid motifs on the Human Metabolome Database**

## Star Methods

### Resource availability

#### Lead Contact

Further information and requests for resources and reagents should be directed to and will be fulfilled by the Lead Contact, James M. Berger.

#### Materials Availability

All reagents generated for this study are available on request through the Lead Contact.

#### Data and Code Availability

The data obtained in this study will be accessible at the NIH Common Fund’s NMDR (supported by NIH grant, U01-DK097430) website, the Metabolomics Workbench, https://www.metabolomicsworkbench.org

### Experimental Model and Subject Details

#### Yeast strains and growth conditions

Crude metabolite samples were extracted from cultures of BY4741 yeast (*MATα his3Δ1 leu2Δ met15Δ0 ura3Δ0*). Growth experiments to assess the sensitivity of yeast to topo II-targeted drugs were performed with a drug-efflux deficient (ED) strain (*pdr1Δ::pdr1:Cyc8 LEU2*) described in (Stepanov et al., 2008). To generate the ED-*mpc1Δ* strain, the KANMX resistance marker was PCR amplified from the pUG6 plasmid (Güldener et al., 1996) with the following primers containing homology regions outside of the *MPC1* gene: (5’)CAGCAAACGTCAATACATCTACATATATACGTATAGATTTTATTGCACTGTGATCGACATGGAGGCCCAGAATACC and (5’)GTTTCCATCTAGTCACCTACTTCAGGTTCTTAGACTGCTCGTTTTACCAGTATAGCGACCAGCATTC. The deletion construct was then transformed into ED yeast. After recovery on YPD plates followed by selection on G418 containing replica plates, *mpc1Δ* mutants were verified by PCR.

CRISPR/Cas9 technology was used to incorporate a 3xHA-tag at the N-terminus and C-terminus of endogenous topo II in the BY4741 background. The following gRNA sequences were integrated in the CRISPR/Cas9 vector, pJH2972 (Anand et al., 2017): (5’)GCAGTGAAAGATAAATGATCGTTGACATGGTTAGCCGTGCGTTTTAGAGCTAGA AATAGC (N-terminal gRNA) and (5’)GCAGTGAAAGATAAATGATCAAAAAGAATGGCGCTTTCTCGTTTTAGAGCTAGA AATAGC (C-terminal gRNA). The CRISPR/Cas9 constructs were then transformed into BY4741 yeast with their corresponding repair templates: (5’)TTTCAGTTAAAGGAGTTTATAACGACGAGCACGCCTAACCATGTACCCATACGATGTTCCTGACTATGCGGGCTATCCCTATGACGTCCCGGACTATGCAGGATCCTATCCATATGACGTTCCAGATTACGCTGGTACGATGTCAACTGAACCGGTAAGCGCCTCTGATAAATATCAGA (N-tag) and (5’)GGAAAACCAAGGATCAGATGTTTCGTTCAATGAAGAAGATGGTACGTACCCATACGATGTTCCTGACTATGCGGGCTATCCCTATGACGTCCCGGACTATGCAGGATCCTATCCATATGACGTTCCAGATTACGCTTGAATAATATTTATCGAGAGAAAGCGCCATTCTTTTTATA (C-tag). Integration of the repair constructs was verified by colony PCR followed by BamHI digest of the product (only the PCR product from a strain with correct integration of the 3xHA-tag will have a BamHI cut site).

Cultures for metabolite extraction were grown in synthetic media in which ammonium sulfate was replaced with monosodium glutamate as the nitrogen source (SC-sulfate). Per liter, SC-sulfate media contains 75 mg of each amino acid, 20 mg of adenine, 75 mg of uracil, 25 mg of inositol, 1.71 g of YNB-sulfate (Sunrise Science), 1 g monosodium glutamate, and 20 g dextrose. Growth assays were conducted in SC media (Sunrise Science). Glucose (20 g/L), lactate (2%) and glycerol (1.5%) were added to the media after autoclaving as required for each experiment.

### Method Details

#### S. cerevisiae metabolite extraction

BY4741 cultures grown in SC-sulfate media were harvested at early log phase, OD_600_ = 0.4-0.8, by vacuum filtration through a surfactant-free cellulose acetate membrane with a 0.2 µm pore size (ThermoFisher Scientific). When ∼5 mL of media was left in the filter, 30 mL of 100 mM NH_4_OAc – TEA (pH 8.0) was added to wash the cells and remove any remaining extracellular material. Immediately after the wash solution was passed through the filter, the filter apparatus was released from vacuum and 10 mL of 90% MeOH, 10 mM NH_4_OAc – TEA (pH 8.0) (chilled to −80 °C) was added to the filter for rapid quenching and extraction (Boer et al., 2010; Crutchfield et al., 2010). The extract was then centrifuged at 4000g for 10–15 min to remove cell debris. The extract supernatant was passed through a 3 kDa MWCO filter (Cytiva) to remove large macromolecules. The filtrate was diluted 2-fold with water and lyophilized to remove the extraction solvent. The remaining extracts were stored at −80°C for up to one month.

#### Supercoil relaxation, decatenation, and ATPase activity assays

Recombinant eukaryotic proteins were prepared as described previously (Lee et al., 2017). Briefly, *Sc*Top2, *Hs*Top2A, and *Hs*Top2B were expressed in the *S. cerevisiae* strain BCY123 (Wasserman and Wang, 1994), and cell pellets were lysed by cryogenic grinding. Proteins were first purified by Ni-affinity purification (HisTrap FF, GE) followed by cation exchange (HiTrap SP, GE). Affinity tags were removed by His_6_-tagged TEV protease (QB3 MacroLab). The digested proteins were passed over a second Ni-affinity column to remove the cleaved His_6_-tag and the His_6_-TEV protease. For the final purification step, proteins were run on a gel-filtration column (S-400, GE). Recombinant *Ec* topoIV was prepared as described previously (Vos et al., 2013). His_6_-tagged subunits of *Ec* topoIV (ParE and ParC) were expressed in BL21 codon-plus (DE3) RIL cells and cells were lysed by sonication. Proteins were first purified by Ni-affinity purification (HiTrap Ni^2+^, GE). After removing the affinity tags by TEV cleavage and a second Ni-affinity step as described above, the proteins were purified by gel filtration (S-300, GE).

Negatively supercoiled pSG483, a derivative of pBluescript SK, was prepared from XL1-Blue *E. coli* cultures by maxi prep as described previously (Lee et al., 2017). Metabolite extracts were solubilized in 12.5% DMSO and metabolite stocks prepared from commercial compounds were solubilized in water and adjusted to pH 6.5-7.5 with HOAc or KOH.

For strand passage activity assays, enzyme stocks were first diluted in two-fold steps in 30 mM Tris-HCl (pH 7.9), 500 mM KOAc, 10% glycerol and 0.5 mM TCEP. *Ec* topo IV subunits were incubated together on ice at high concentrations (200 - 300 µM of each subunit) for 10 min prior to dilution to allow formation of the holoenzyme. The enzyme was then added to the appropriate DNA substrate, negatively supercoiled pSG483 plasmid for supercoil relaxation assays or kDNA (Inspiralis) for decatenation assays, followed by metabolites. Reactions were initiated by the addition of ATP (5 mM for assays with extracts and 1 mM for assays with commercial metabolites) and shifted to 30 °C for *Sc*Top2 or 37 °C for *Hs*Top2A/B and topo IV. The final reaction conditions contained 32 mM Tris-HCl (pH 7.9), 100 mM KOAc, 20 mM Mg(OAc)_2_, 0.05 mg/ml BSA, 0.6 mM TCEP, and 10% glycerol. Assays with metabolite extracts also contained 2.5% DMSO. Decatenation reactions were incubated for 3 min and supercoil relaxation assays were incubated for 5 min. Reactions were quenched with 20 mM EDTA and 1% SDS and treated with 0.2mg/mL Proteinase K to remove any remaining protein bound to DNA. Final products were resolved and visualized by native gel electrophoresis (1.4% agarose, 1X TAE) run at 2-2.5 V/cm for 15-20h. Supercoil relaxation products were run on native gels and decatenation products were run on gels containing 0.4 μg/mL ethidium bromide.

ATP hydrolysis by topo II was measured by an NADH-coupled ATPase assay. Reactions were prepared with 50 nM *Sc*Top2, 200 ng/µL sheared salmon sperm DNA (Fisher Scientific), 0.5 mM NADH, 0.2% rabbit muscle pyruvate kinase/lactate dehydrogenase (Sigma-Aldrich), 2 mM PEP, 36 mM Tris-HCl (pH 7.9), 100 mM KOAc, 10 mM Mg(OAc)_2_, 0.05 mg/mL BSA, 0.6 mM TCEP, and 2% glycerol in a final reaction volume of 75 µL. ATP concentrations were titrated from 0-4 mM. Depletion of NADH was observed over the course of one hour at 30°C by monitoring absorbance at 340 nm in a CLARIOStar microplate reader (BMB LAB TECH). ATP hydrolysis rates were calculated based on an NADH standard curve and fit for K_m_ and V_max_ values in PRISM (GraphPad).

#### Fractionation and purification of active compounds from crude metabolite extracts

Lyophilized samples of metabolite extracts were resuspended with water and adjusted to pH 3.5 with formic acid or pH 10 with TEA. The metabolite solution was transferred to a separatory funnel and mixed with an equal volume of butanol. The mixture was then left until the separation of aqueous and organic layers was complete. The aqueous solvent was removed by lyophilization and the organic solvent was removed in a rotary evaporator. The activities of the aqueous and organic fractions from acid- or base-extraction were assessed in a supercoil relaxation assay to determine the hydrophilic nature of the active compound.

Enrichment and purification of the stimulatory metabolite from crude extracts for LC-MS/MS analysis began with solid phase extraction (SPE). Crude extracts were solubilized in water and adjusted to pH 10 with TEA. A mixed-mode anion exchange cartridge was equilibrated as per the manufacturer’s instructions (Oasis MAX, Waters). The flow-through from the passage of the crude extract through the SPE column (i.e., unbound material) was lyophilized for further purification.

The extract was then further purified by reverse-phase HPLC on a C18 column (Atlantis T3, Waters). One column volume of water with 0.1% formic acid followed by eight column volumes of MeOH were passed through the column over the course of the HPLC run. Absorbance at 260 nm and 280 nm wavelengths was observed by a Waters 2996 Photodiode array. Fractions from the aqueous phase were lyophilized and assessed for stimulation of supercoil relaxation. Active fractions were then further purified through normal-phase HPLC on an amide column (XBridge BEH Amide, Waters). One column volume of 80% acetonitrile with 0.1% formic acid followed by a three-column volume gradient down to 40% acetonitrile with 0.1% formic acid were passed through the column. Afterwards, one column volume of water with 0.1% formic acid was passed over the column as the last step. The separated fractions were lyophilized and assessed similarly to all prior purification steps.

#### Antarctic phosphatase and phosphodiesterase treatment of metabolites

Snake venom phosphodiesterase I (SVPD, Sigma-Aldrich) was reconstituted at 1 mg/mL in 20 mM Tris-HCl (pH 7.9), 100 mM NH_4_OAc, 20 mM Mg(OAc)_2_, 50% glycerol (Konokhova et al., 2016). Crude extracts were first fractionated by SPE as described above. The lyophilized sample was then resuspended in water and split into four samples of 44 μL each. Next, 5μl of 10X reaction buffer was added to each sample: 10x Antarctic phosphatase (AP) reaction buffer (New England BioLabs) was added to two samples and 200 mM Tris-HCl (pH 7.9), 1 M NaCl, 200 mM MgCl_2_ was added to the remaining two samples for SVPD treatment. For each pair of samples, 1 μl of water was added to the no enzyme control and 1 μl of enzyme (AP or SVPD) was added to the second sample. All reactions were incubated at 37 °C for two hours. Samples were then diluted to a 1 mL final volume and passed through a 3 kDa MWCO filter. The filtrate was lyophilized and assessed for activity against topo II in a supercoil relaxation assay.

#### Untargeted LC-MS/MS analysis of metabolite samples

Lyophilized samples were reconstituted in water plus 0.1% (v/v) formic acid such that samples reached a final concentration of 20 mg/ml. A pooled sample was generated by combining equal volume aliquots from each sample. Untargeted metabolomics uHPLC-MS/MS data acquisition was performed in both positive and negative ion modes and using both reversed-phase and normal-phase separations.

Liquid chromatographic separation was performed using a Dionex Ultimate 3000 uHPLC system (Thermo) for reversed-phase and normal-phase methods. For both methods, mobile phase A was 1mM ammonium acetate + 0.1% (v/v) formic acid in water and mobile phase B was 1mM ammonium acetate + 0.1% (v/v) formic acid in 20% water / 80% acetonitrile (v/v). Reversed-phase experiments were performed with a Hypersil GOLD aQ polar endcapped C18 column (150 mm x 2.1 mm, 1.9 µm particle size) (Thermo) using the following gradient of mobile phases: 0% B from 0-2 min, 0-60% B from 2-13 min, 60-90% B from 13-15 min, 90-0% B from 15-16 min, and 0% B from 16-20 minutes. Normal-phase experiments were performed with a Polyhydroxyethyl A hydrophilic interaction chromatography column (200 mm x 2.1 mm, 5 µm particle size) (PolyLC) using the following gradient of mobile phases: 80% B from 0-2 min, 80-40% B from 2-13 min, 40-10% B from 13-15 min, 10-80% B from 15-16 min, and 80% B from 16-20 min. For both uHPLC methods, the flow rate was set to 0.2 mL/min. Samples were each injected 3 times under each unique set of conditions. A blank injection of 0.1% formic acid in water (v/v) was run in between each sample injection to minimize carry over.

After separation by uHPLC, metabolites were ionized and detected by a Q-Exactive quadrupole-orbitrap mass spectrometer (Thermo) using a heated electrospray ionization source. Samples were run separately in negative and positive ion modes. In positive mode, ionization was performed with a spray voltage of 4000 V, capillary temperature of 275°C, sheath gas flow rate of 35 Arb, and auxiliary gas flow rate of 8 Arb. Negative mode ionization was performed with a spray voltage of 4000 V, capillary temperature of 350°C, sheath gas flow rate of 25 Arb, and auxiliary gas flow rate of 2 Arb.

For each set of separation and ion modes, mass spectra were acquired by scans in data-dependent MS/MS mode for the pooled samples and full-MS mode for individual fraction replicates. Full-MS scans were performed at a resolution of 140,000 and a mass range of m/z 65-850. Data-dependent MS/MS scans were carried out for the top-5 abundant ions at a resolution of 140,000 FWHM with a dynamic exclusion of 6.0 seconds and stepped normalized collision energy of 20, 40, and 100. MS/MS fragments were observed at a resolution of 17,500 FWHM.

The LC-MS and LC-MS/MS data were processed using Compound Discoverer 3.1 software (ThermoFisher Scientific). Spectral features retention times were aligned with a permitted mass deviation of 5 ppm. Background peaks were subtracted based on features in a blank sample. The data-dependent MS/MS files were analyzed for compound identification using mzCloud and the ChemSpider yeast metabolome database. The full-MS scan data was used for comparison of metabolite abundance by average peak area. Analysis of metabolite enrichment was performed by comparing replicates from fractions with an effect on topo II activity to those with no effect on topo II activity.

#### Targeted Metabolomics Analysis of TCA Metabolites

Lyophilized samples of crude extracts from yeast grown in different nutrient conditions were reconstituted in water + 0.1% formic acid, such that the ratio of milligrams of cell mass to microliters of acidified water was 0.5. Samples were analyzed on a Dionex Ultimate 3000 uHPLC (Thermo) coupled to a TSQ Vantage triple quadrupole mass spectrometer (Thermo). Analytes were separated on a Hypersil GOLD aQ polar endcapped C18 column (150 mm x 2.1 mm, 1.9 µm particle size) using an isocratic flow of 100% mobile phase A (water +0.1% formic acid) at a flow rate of 0.2 mL/min over five minutes. Analytes were ionized by heated electrospray ionization with a spray voltage of 3000 V, capillary temperature of 204°C, sheath gas flow rate of 50 Arb, and auxiliary gas flow rate of 55 Arb. Metabolites were detected in negative ion mode using distinct single reaction monitoring scans with the following transitions and collision energies - citric acid: m/z 191 → 111 (CE 12), alpha-ketoglutaric acid: m/z 145 → 101 (CE 10), succinic acid: 117 → 73 (CE 12), fumaric acid: 115 → 71 (CE 10), malic acid: 133 → 71 (CE 15). Chromatographic peak areas were used for comparisons of metabolite abundance. Data were normalized to the average of the control condition and unpaired, two-tailed t-tests were performed in PRISM (GraphPad) to determine significant differences.

#### S. cerevisiae growth assays

Stock solutions of etoposide (Sigma-Aldrich) and ICRF-187 (TCI) were prepared at 100 mM concentration in 100% DMSO then aliquoted and stored at −20 °C. DMSO was added to all no drug experimental controls to match the DMSO content of the drug containing conditions. For spot growth assays, ED yeast were first grown to saturation at 30 °C overnight in SC media. The starter culture was then diluted to OD_600_ = 0.1 in sterile water. In a sterile 96-well plate, the culture was serially diluted down from OD_600_ = 0.1 in 5-fold dilution steps. A multichannel pipette was then used to spot 5 μL of each dilution on an SC agar plate with the appropriate drug condition for each experiment.

Plates were incubated for 2-3 days at 30 °C and imaged by normal photography. For liquid media growth assays, ED or ED-*mpc1Δ* yeast were first grown to saturation at 30°C overnight in SC media with the appropriate carbon source for each experiment. The cultures were then diluted to OD_600_ 0.01 in the growth assay media condition and placed in a sterile 24-well plate with 1.5ml of culture in each well. The plates were incubated at 30 °C with constant shaking inside a Bio-Tek Synergy HT plate reader for 30 - 35 h. OD_600_ measurements were taken every 15 min to generate a growth curve for each well.

#### Western Blot

Approximately 1.5 x 10^8^ cells or 15 OD_600_ units of log phase yeast (OD_600_ = 0.8-1.2) were harvested by centrifugation. Pellets were then resuspended in 200 μL of 10% trichloroacetic acid (TCA) and incubated at room temperature for 30 min. The TCA solution was then removed by centrifugation and the pellets were washed with 1 mL of 1 M HEPES•KOH (pH 7.5). After removal of the wash solution by centrifugation, the pellets were then resuspended in 50 μL 2x SDS-PAGE loading buffer with ∼50 μL of 0.5 mM glass beads and vortexed for 3 min. An additional 50 μL of 2x SDS-PAGE loading buffer was added before the samples were boiled for 5 min and then vortexed for 15 sec. After centrifugation to pellet the beads, 20 μL of each supernatant sample were run on an SDS-PAGE gradient gel and transferred onto a PVDF membrane. The membranes were blotted with rat anti-HA (1:5000, Roche 11867423001) and rabbit anti-tubulin (1:5000, abcam ab184970) followed by IRDye 800CW goat anti-Rat (1:15,000, Li-Cor 926-32219) and IRDye 680CW goat anti-rabbit (1:15,000, Li-Cor 926-68071). Blots were imaged on a Li-Cor Odyssey system and band intensities were analyzed in ImageJ.

### Quantification and Statistical Analysis

Data are presented as the mean ± SD of a minimum of three independent experiments, indicated by the n value described in the figure legends.

